# Unlocking the multiphasic nature of intracellular calcium signatures triggered by fungal signals in *Lotus japonicus* roots

**DOI:** 10.1101/2022.04.13.487819

**Authors:** Filippo Binci, Elisabetta Offer, Andrea Crosino, Ivan Sciascia, Jürgen Kleine-Vehn, Andrea Genre, Marco Giovannetti, Lorella Navazio

## Abstract

The recognition of different microbe-associated molecular patterns in the rhizosphere triggers in the plant root the activation of either an immune response or an accommodation program. In both types of responses, Ca^2+^ is a crucial intracellular messenger, mediating the early stages of the respective signalling pathways. In this work, we analysed the cytosolic and nuclear Ca^2+^ changes activated by a set of chitin-related oligomers in different genetic backgrounds of *Lotus japonicus* roots by using specifically targeted aequorin-based Ca^2+^ reporters. By means of pharmacological and genetic approaches, we dissected the Ca^2+^ signal into two temporally distinct components: a rapid initial transient, followed by a longer and milder elevation in Ca^2+^ concentration. Taking advantage of a complementary analysis using a cameleon-based bioassay in *Medicago truncatula* root organ cultures, we showed that the second phase can be interpreted as the Ca^2+^ spiking that is widely described in response to the perception of symbiotic signals. By contrast, the rapid first phase, critically dependent on elicitor concentration, was found to correlate with the activation of plant immunity marker genes. Overall, our study provides novel clues to a better understanding of the subtle boundaries between symbiotic and immunity responses in root-fungus interactions.

**Highlight:** Intracellular calcium changes induced in *Lotus japonicus* roots by fungal signals were dissected in two separate phases, relying on distinct genetic programs and differentially mediating plant symbiotic or immunity responses.

## Introduction

In the rhizosphere, plant roots encounter myriads of microorganisms that can activate distinct transcriptional and developmental programs upon the recognition of microbe-associated molecular patterns (Verbon and Liberman, 2016; Bonhomme *et al*., 2021; Delaux and Schornack, 2021; Chiu *et al*., 2022). These include highly conserved microbial components such as chitin or flagellin, normally driving the plant response towards an immunity-like program (Wan *et al*. 2012; Platre *et al*., 2022), as well as specific microbe-released signals such as Myc-factors and Nod-factors, two classes of diffusible chitin-based molecules that trigger root endosymbiotic programs, in arbuscular mycorrhizas and legume nodulation, respectively (Choi *et al*., 2018; Ghahremani and MacLean, 2021). In both cases - immunity and symbiosis -, although microbial signals are perceived by the plant host through a set of plasma membrane-associated receptors (Zipfel and Oldroyd, 2017), a pivotal common mediator of most downstream signalling pathways is calcium. Indeed, transient increases in intracellular calcium concentration ([Ca^2+^]) have been shown to be generated during plant responses to both pathogens (Ranf *et al*., 2011; Wan *et al*., 2012; Nars *et al*., 2013; Keinath *et al*., 2015; Feng *et al*., 2019; Zhang *et al*., 2021, Köster *et al*., 2022) and symbionts (Navazio *et al*.,2007; Capoen *et al*., 2011; Genre *et al*., 2013; Feng *et al*., 2019; Zhang *et al*., 2021). Such Ca^2+^-mediated signals display subtle and not fully elucidated differences (the so-called Ca^2+^ signature) that are believed to control the activation of tailored downstream molecular, cellular and metabolic responses (Zipfel and Oldroyd, 2017). In this frame, crosstalks and overlaps have been highlighted in plant symbiotic and pathogenic interactions between microbe-released molecules, receptor roles and Ca^2+^-mediated signal transduction pathways (Ried *et al*., 2019; Zhang *et al*., 2021).

In this research, we focused on arbuscular mycorrhizal (AM) symbiosis, the most ancient and widespread root endosymbiosis established between the vast majority of land plants and Glomeromycotina fungi (Choi *et al*., 2018; Genre *et al*., 2020). AM fungi-released Myc-factors include short-chain chitin oligomers, such as tetrameric chitooligosaccharides (CO4), and lipochitooligosaccharides (mycLCOs) with structural similarities to rhizobial Nod-factors (Maillet *et al*., 2011; Genre *et al*., 2013; Feng *et al*., 2019). The recognition of Nod- and Myc-factors and the activation of the respective symbiotic programs rely on the Common Symbiotic Signalling Pathway (CSSP), encompassing a diverse set of co-receptors (LjSYMRK/MtDMI2), cation channels (LjCASTOR/MtDMI1, LjPOLLUX, MtCNGC15), Ca^2+^ transporters (MtMCA8) and Ca^2+^ sensor proteins (LjCCaMK/MtDMI3) (Charpentier, 2018; Choi *et al*., 2018; Radhakrishnan *et al*., 2020). Nevertheless, the cell wall of AM fungi - like all fungi - contains a large amount of long-chain chitin, a well known pathogen-associated molecular pattern (PAMP) eliciting immunity-related responses in plants. For this reason, a number of recent studies has investigated the role of different chitin-based molecules in AM signalling, with sometimes contrasting results, likely depending on the experimental setup. While, for example, short-chain (but not long-chain) chitooligosaccharides were shown to trigger AM-specific Ca^2+^-based symbiotic signals (Genre *et al*., 2013), a different study suggested that long-chain oligomers such as CO8 trigger both pathogenic and symbiotic signalling (Feng *et al*., 2019). This ambiguity is also due to our limited understanding of the complexity of signal exchange during early plant-microbe interactions and the role of plant receptors and co-receptors in different interactions and different plant species (Yu *et al*., 2017; Zhang *et al*., 2021). In fact, the rice chitin receptor *OsCERK1* is crucial for both AM symbiosis and immunity (Zhang *et al*., 2014, Miyata *et al*., 2014, Carotenuto *et al*., 2017) and the discrimination between the two responses has recently been shown to depend on the competition between two alternative co-receptors for associating with *Os*CERK1 in the presence of different ligands (Zhang *et al*., 2021). The scenario is even more complex in legumes, where the family of LysM receptor-like kinases (which include the receptors for chitin-based signals) is much more expanded than in other plants. In *Lotus japonicus, LjCERK6* is responsible for chitin-triggered immunity in response to fungal pathogens (Bozsoki *et al*., 2017) but the corresponding receptors involved in AM symbiosis remain elusive (Chiu *et al*., 2020). Altogether, a picture is emerging where the composition of fungal signals and the overlapping roles of plant plasma membrane receptors generate an intricate continuum between immunity and symbiosis (Zhang *et al*., 2021).

A second level of confounding factors derives from the use of distinct methodological approaches to the analysis of Ca^2+^-mediated plant responses (Costa *et al*., 2018; Grenzi *et al*., 2021). On the one hand, intracellular Ca^2+^ elevations in response to the perception of different PAMPs have mainly been mainly investigated in entire plants by using the cytosolic-expressed bioluminescent probe aequorin (Ranf *et al*., 2011; Monaghan *et al*., 2015). On the other hand, CO- and LCO-triggered oscillations in perinuclear and nuclear Ca^2+^ concentrations (the so-called Ca^2+^spiking) have been characterised at the single-cell level with the use of fluorescent genetically encoded calcium indicators (GECIs), such as cameleon or GECO (Genre *et al*., 2013; Kelner *et al*., 2018; Feng *et al*., 2019; Zhang *et al*., 2021). This has largely hampered the possibility of a direct comparison between the two sets of results, with few exceptions. Flagellin- and chitin-induced cytosolic Ca^2+^ elevations revealed an oscillatory nature when analysed with Yellow Cameleon 3.6 and GECO (Thor and Peiter *et al*., 2014, Keinath *et al*., 2015). Furthermore, symbiotic signals were found to induce a characteristic cytosolic Ca^2+^ elevation (Miwa *et al*., 2006; Navazio *et al*., 2007) that was described as a Ca^2+^ influx and - in the case of legume nodulation - was required for symbiosis development (Morieri *et al*., 2013). In short, the need is emerging to investigate and integrate Ca^2+^-mediated plant responses to biotic signals by using multiple approaches and combining qualitative (Ca^2+^ imaging) and quantitative (Ca^2+^ measurement) approaches.

In this work, we analysed the effects induced by a set of chitin-related oligomers on cytosolic and nuclear Ca^2+^ levels in different genetic backgrounds of *L. japonicus* roots by using specifically targeted aequorin-based Ca^2+^ reporters. We accurately quantified the root Ca^2+^ signatures upon treatment with specific fungal molecules and showed that CO4-, CO8- and mycLCO-induced Ca^2+^ elevations are distinct for their intracellular localization, temporal dynamics and plant genetic requirements. An alternative approach based on Ca^2+^ imaging with a nuclear-localized cameleon probe in *Medicago truncatula* root organ cultures further supported our description of two distinct temporal phases in the observed Ca^2+^ responses, depending on the concentration of the stimulus and the genetic background. Our analyses of the overall plant root response in terms of intracellular Ca^2+^ changes help disentangling the complex communication circuits in the rhizosphere and enhance our quantitative understanding of how plant roots can perceive and transduce different fungal molecules.

## Materials and Methods

### Molecular cloning and bacterial transformation

The nucleotide sequence encoding the bioluminescent Ca^2+^ indicator aequorin fused to Yellow Fluorescent Protein (YFP) and targeted to either the cytosol-only (CPK17_G2A_-NES-YA) or nucleus-only (NLS-YA) (Mehlmer *et al*., 2012) were amplified with Q5 DNA polymerase (NEB) according to manufacturer’s instructions. The promoter (1100 bp upstream of the ATG) of ubiquitin10 was amplified from the genome of *Lotus japonicus* (*Gifu* ecotype). Primers (listed in Table S1) were designed to be compatible with the GreenGate cloning system and cloning of the entry vectors was performed according to it (Lampropoulos *et al*., 2013). Expression vectors were assembled via cut-ligation with BsaI (NEB) and T4-DNA ligase (NEB), following the GreenGate protocol. The selection and amplification of vectors were performed in DH5α *E. coli* cells. The sequences were checked via Sanger sequencing at BMR Genomics (Padova, Italy) and by Primordium long-read DNA sequencing at Primordium Labs (Arcadia, CA, USA). The expression vectors were then transformed into *Agrobacterium rhizogenes 1193* via the freeze & thaw method (Wise *et al*., 2006). Expression vectors are listed in Table S2.

### Generation of Lotus japonicus composite plants

*L. japonicus* ecotype Gifu seeds were scarified with sandpaper and sterilized in 0.5% (w/v) sodium hypochlorite for 11 min. Seeds were then rinsed and washed 5 times in sterile distilled water. For germination, seeds were placed into sterile Petri dishes with H_2_O with 1% (w/v) plant agar and wrapped in aluminum foil. After 3 days at 23°C, young seedlings were transferred to square plates (12×12 cm) containing ½ strength B5 growth medium, supplemented with 0.8% (w/v) plant agar and adjusted to pH 5.5 with 1 M KOH. Plates were grown vertically under long-day conditions (23°C, 16 h light/8 h dark cycle). After 3 days, the seedlings were transformed via *A. rhizogenes 1193* - mediated hairy root transformation (Boisson-Dernier *et al*., 2001). The root was cut off with a blade and the wound was dipped into a fully grown plate of *A. rhizogenes* carrying the plasmid of interest. The infected shoots were then placed onto square plates (12×12) containing ½ strength B5 growth medium, 0.8% plant agar, pH 5.5. After 16 h in the darkness, seedlings were co-cultivated with bacteria for 3 days. Afterward, the seedlings were transferred into new squared plates with the same medium supplemented with 300 mg/ml cefotaxime and 0.1% Plant Preservative Mixture (PPM, Duchefa) under long-day conditions (23°C, 16 h light/8 h dark cycle). After 3 weeks, transformed roots were checked for expression of the transformation marker (pAtUBQ10::mCherry) using the stereomicroscope MZ16F (Leica). The intracellular localization of the probe was confirmed by confocal microscopy observations (Zeiss LSM900 Airyscan2). Transformed plants were kept in the same growth medium under long-day conditions (23°C, 16 h light/8 h dark cycle).

### Aequorin-based Ca^2+^ measurement assays

For Ca^2+^ measurement assays in *L. japonicus* composite plants, 5 mm segments of transformed roots expressing aequorin targeted to either the cytosol or nucleus were reconstituted overnight with 5 mM coelenterazine. On the following day, after extensive washing, each root piece was placed in the dark chamber of a custom-built luminometer (Electron Tubes) containing a 9893/350A photomultiplier (Thorn EMI). The root was placed in 50 μl H_2_O and challenged by injection of an equal volume of a 2-fold concentrated solution for each tested stimulus: CO4, CO8 (IsoSep), mycLCOs (equimolar mix of both non-sulfated and sulfated C16:0 and C18:1). The stock solutions were 10^−3^ M in 50% ethanol for CO4, 10^−4^ M in 50% ethanol for CO8, 10^−3^ M in DMSO for mycLCOs. Controls were performed by injecting an equal volume of the solvents in which the compounds are dissolved at the working concentration. Ca^2+^ dynamics were recorded for a total of 30 min before the injection of 100 μl of the discharge solution (30%, v/v, ethanol, 1 M CaCl2). The light signal was collected and converted offline into Ca^2+^ concentration values using a computer algorithm based on the Ca^2+^ response curve of aequorin (Brini *et al*., 1995). All Ca^2+^ concentration values for each biological replicate are available (Table S3). For pharmacological analyses, the root pieces, before challenge with chito-oligomers, were pre-treated with the pharmacological agents for different time intervals: 1.5 mM LaCl_3_ (Sigma) and 2 mM EGTA (Sigma) for 10 min, 50 μM cyclopiazonic acid (CPA, stock solution 30 mM in DMSO) for 1 h, 10 μM VAC1 (stock solution 10^−3^ M in DMSO) for 2.5 h (Dünser *et al*., 2021).

### Cameleon-based Ca^2+^ imaging assays

*Medicago truncatula* genotype Jemalong A17 and the *dmi2-2* mutant (Catoira *et al*., 2000) were genetically transformed using *A. rhizogenes* to generate root organ cultures expressing the nuclear-localized 35S:NupYC2.1 cameleon construct (Sieberer *et al*., 2009), as described in Chabaud *et al*. (2011). Ca^2+^ imaging was conducted on excised young lateral roots, placed in 2mm-thick microscope slide microchambers. The water in the microchamber was rapidly (< 30 s) substituted by 200 μl of CO4 solution before initiating confocal image acquisition within 1-3 minutes. FRET-based ratio imaging of the YFP and CFP cameleon fluorescence was used for the detection and plotting of relative changes in nuclear Ca^2+^ levels (Chabaud *et al*., 2011).

### Gene expression analysis

*L. japonicus Gifu* seeds were sterilized as described above and the seedlings were grown for 10 days (23°C, 16 h light/8 h dark cycle) in vertical square plates (12×12 cm) containing ½ strength B5 growth medium, 0.8% (w/v) agar, pH 5.5. Groups of 12 plants were treated in 4 ml solutions containing either 10^−7^ M CO4, 10^−9^ M CO4 or the control treatment (50% ethanol diluted 1:1000). The grouped samples were harvested after 1 h of treatment. During harvesting, the roots were cut from the shoots and immediately frozen in liquid N_2_ in a 2 ml tube containing two metal beads. The root material was homogenized with TissueLyser II (Qiagen) at 30 Hz for 45 seconds. Total RNA was extracted using the RNeasy Plant Mini Kit (Qiagen) following the manufacturer’s instructions. RNA quantity and quality were checked by nanodrop and agarose gel electrophoresis. After DNAse I (Invitrogen) treatment, cDNA synthesis was performed with random hexamers using the RevertAid RT Kit (Thermofisher). qRT-PCR was performed using the HOT FIREPOL EvaGreen qPCR mix plus (Solis Biodyne) on a 7500 real-time PCR system (Applied Biosystems). The *LjUbiquitin10* gene was used as an internal reference for analysis of the target gene expression. All the primers (Table S1) used for the qRT-PCR analyses have been previously published (Nakagawa *et al*., 2011; Giovannetti *et al*., 2015; Bozsoki *et al*., 2017).

### Statistical analysis and data visualisation

Data were statistically analysed and presented graphically using R statistical environment and Rstudio (RStudio Team, 2020; R Core Team, 2022). When passing its assumptions, the Anova test and Tukey’s post-hoc test was applied, in the other cases, Kruskal Wallis and Dunn’s tests were performed (Table S3). R scripts (Supplementary Dataset S1) and raw data (Supplementary Dataset S2-S8) ensure full reproducibility of statistical analysis and plots.

## Results

### Different fungal signals activate apparently similar Ca^2+^ transients in both the cytosol and nucleus of L. japonicus roots

To monitor intracellular Ca^2+^ dynamics in *L. japonicus* roots, we assembled expression cassettes for YFP-tagged aequorin reporters specifically targeted to either the cytosol or the nucleus (Mehlmer *et al*., 2012) under the control of the *LjUBI10* promoter (Supplementary Fig. S1). The constructs were inserted into *L. japonicus* Gifu background via *A. rhizogenes-mediated* hairy roots transformation and the correct localization of the Ca^2+^ probes was confirmed by confocal microscopy analyses (Supplementary Fig. S1). Root segments from *L. japonicus* composite plants were then challenged with the purified fungal signals CO4 (short-chain COs), CO8 (long-chain COs) and mycLCOs (sulfated and non-sulfated lipo-COs mixture), and the changes in intracellular [Ca^2+^] were measured. All categories of chitin-based oligomers were found to trigger stimulus-specific cytosolic Ca^2+^ signatures, varying according to the intensity and timing of the Ca^2+^ increase. In particular, CO4 and CO8 transiently induced cytosolic Ca^2+^ transients characterised by the highest magnitude of the peak (Fig. 1A-C, Table S3 and Dataset S2). No Ca^2+^ changes were recorded upon administration of solvent controls (Fig. 1A and Dataset S2).

**Fig. 1.**
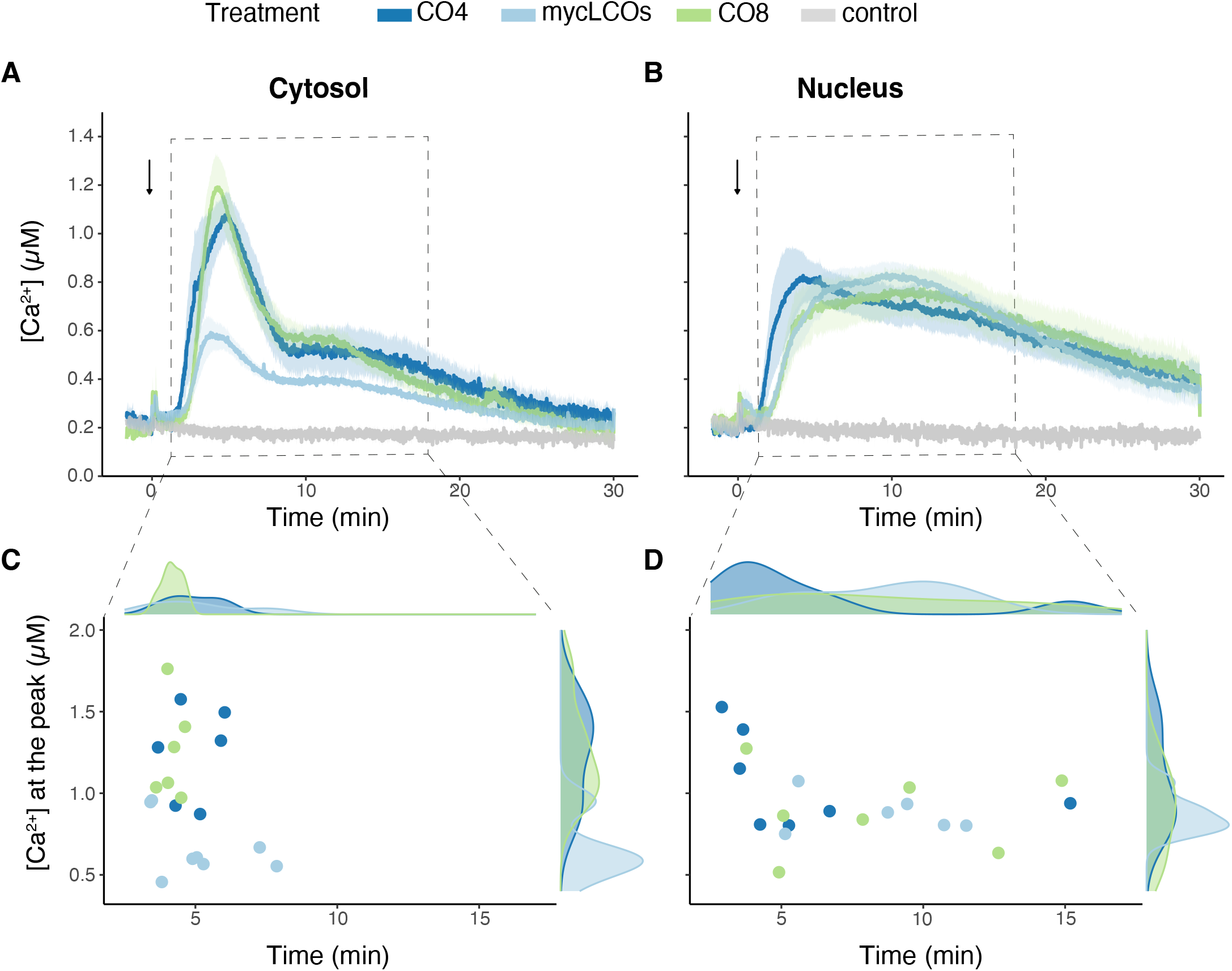
Monitoring of cytosolic and nuclear free [Ca^2+^] in 5 mm-long root segments from composite plants of *L. japonicus* in response to different chitin-based oligomers. *L. japonicus* roots were transformed via *A. rhizogenes* with constructs encoding aequorin chimeras targeted to either the cytosol (A,C) or nucleus (B,D) and subcellular Ca^2+^ dynamics were measured after challenge with 10^−7^ M CO4 (dark blue), 10^−6^ M CO8 (green), 10^−7^ M mycLCOs (light blue) or with the same concentration of the solvents in which the compounds were dissolved (grey). In A-B, data are presented as means ± SE (shading) of n≥6 traces from at least 3 different composite plants (independent transformations). The arrow indicates the time of stimulus injection (time 0). In C-D, dots represent the maximum [Ca^2+^] value for each trace in the time range of 1-18 minutes. The frequency distributions of the dots are represented on top of the panel (time) and at the right of the panel ([Ca^2+^]).

Notably, the overall shape of the observed cytosolic Ca^2+^ dynamics is markedly biphasic, with an initial major peak recorded in the first 8 minutes after stimulation, followed by a broader shoulder. Such dynamics closely resembles those triggered by germinating spore exudates of *Gigaspora margarita* (Navazio *et al*., 2007; Moscatiello *et al*., 2018). CO4, mycLCOs, and CO8 were also found to induce Ca^2+^ transients in the nucleus (Fig. 1B,D), in agreement with the activation of nuclear Ca^2+^ responses by short-chain (Genre *et al*., 2013; Sun *et al*., 2015) and long-chain COs (Feng *et al*., 2019) monitored with cameleon probes. Interestingly, no significant differences could be detected in either the overall shape or peak intensity of the nuclear-induced Ca^2+^ transients triggered by the three categories of microbial signals (Fig. 1 and Table S3).

Overall, the Ca^2+^ transients activated by the different fungal signals, and monitored via aequorin-based reporters, looked apparently similar, apart from the intensity of the early cytosolic Ca^2+^ peak.

### Origin and modulation of the CO4-induced intracellular Ca^2+^ fluxes

To elucidate the source of the observed nuclear and cytosolic Ca^2+^ fluxes activated by chitin oligomers we performed a pharmacological analysis, focusing on CO4, *i.e*. the symbiotic signals that triggered the strongest [Ca^2+^] changes in our experimental conditions. Firstly, in order to assess the contribution of extracellular Ca^2+^ to the generation of the observed Ca^2+^ responses, we pre-treated roots with either the Ca^2+^ channel inhibitor LaCl_3_ or the extracellular chelator EGTA. LaCl_3_ treatment caused a complete abolishment of the Ca^2+^ response to CO4 in both the cytosol (Fig. 2A,C, Table S3 and Supplementary Dataset S3) and nucleus (Fig. 2B,D and Table S3). The cytosolic Ca^2+^ trace was almost completely flattened also in the presence of EGTA, where the main peak of the first phase (within 8 min after the stimulus, hereafter called phase 1) was replaced by a very limited elevation (Fig. 2A,C and Table S3). EGTA only abolished the second part of the Ca^2+^ transient (8-30 min after the stimulus, hereafter phase 2) in the nucleus, while the phase 1 peak was maintained (Fig. 2B,D and Table S3). These experiments suggested that Ca^2+^ in the external milieu can have a more relevant impact on phase 1 Ca^2+^ cytosolic signalling, rather than on the phase 1 peak in nuclear [Ca^2+^].

**Fig. 2.**
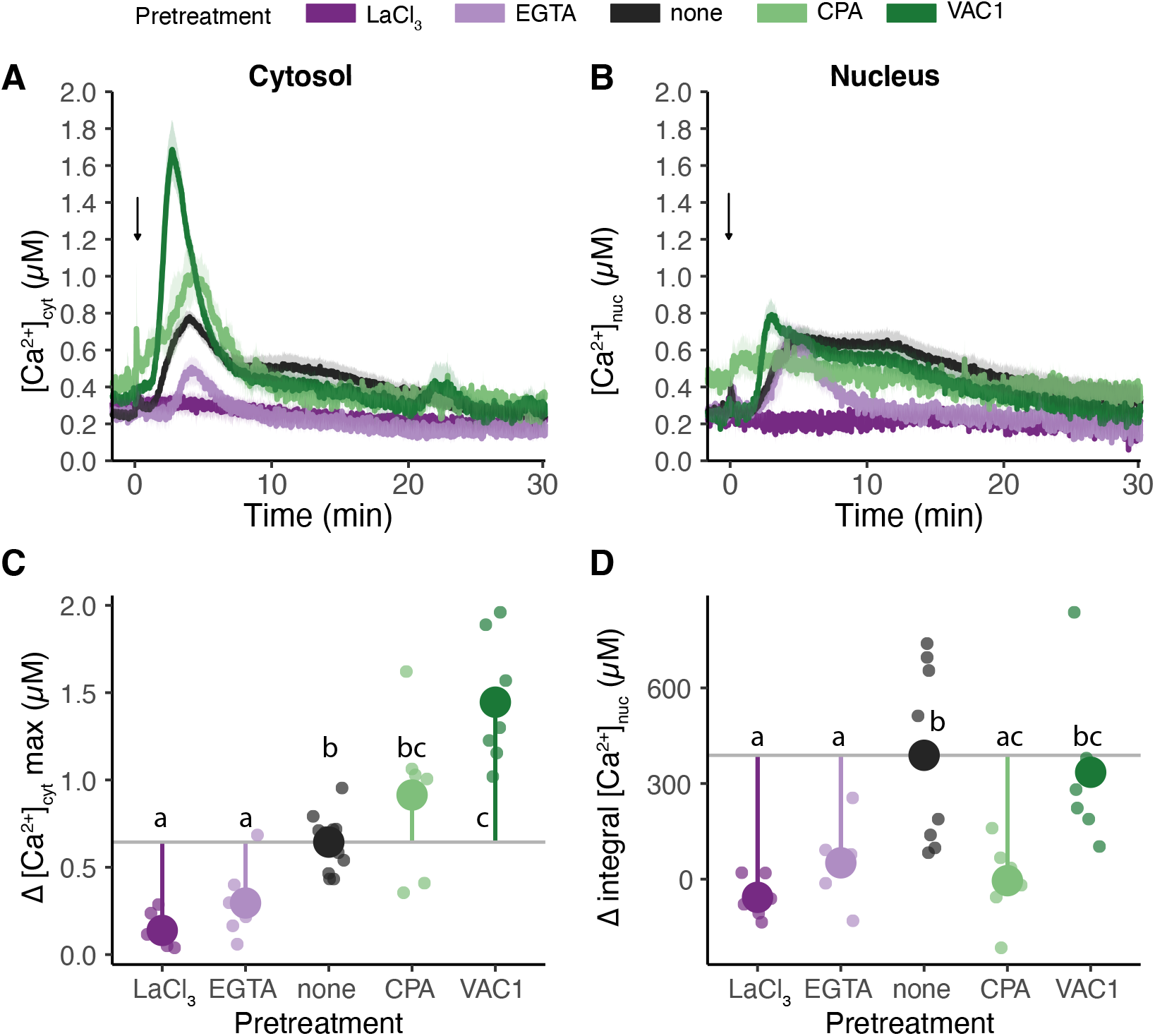
Pharmacological analyses of short-chain COs-induced intracellular Ca^2+^ fluxes in *L. japonicus* roots. Cytosolic [Ca^2+^] (A,C) and nuclear [Ca^2+^] (B,D) dynamics were monitored in 5-mm long root segments from composite plants challenged with 10^−7^ M CO4 after pre-treatment with either 1.5 mM LaCl_3_ (dark violet), 2 mM EGTA (light violet), 50 μM CPA (dark green), 10 μM VAC1 (light green) or none (grey, either H_2_O or solvent control). In A-B, data are presented as means ± SE (shading) of n≥6 traces obtained from at least 3 different composite plants (independent transformation). In C-D, small dots represent the delta maximum cytosolic [Ca^2+^] (Δ[Ca^2+^]_cyt_) (C) and the delta integrated [Ca^2+^] (Δ[Ca^2+^]_nuc_) (D) for each trace in the time range of 2-20 minutes, while the big circles represent the mean. Δ[Ca^2+^] was calculated by subtracting the mean of the resting [Ca^2+^] in the pre-stimulus phase to each [Ca^2+^] measurement value following the stimulus injection (arrow). The coloured vertical line shows the difference between the mean of each treatment and the control (horizontal grey line). Different letters indicate statistically significant differences among groups, according to Kruskal-Wallis test followed by Dunn’s post-hoc correction (p-value <0.05).

We then misregulated Ca^2+^ homeostasis in the plant cell endomembrane system, by applying drug treatments targeting the endoplasmic reticulum (ER) and the vacuole. The first drug we tested was cyclopiazonic acid (CPA), an inhibitor of ER-type Ca^2+^ ATPases present in the ER membrane, causing Ca^2+^ depletion of the ER lumen (De Vriese *et al*., 2018; Cortese *et al*., 2022). Pre-treatment with CPA did not alter the phase 1 peak in cytosolic Ca^2+^ elevation evoked by CO4 (Fig. 2A,C and Table S3), suggesting that the ER to be acting as a Ca^2+^ sink, rather than a source, in this phase. By contrast, CPA completely abolished the Ca^2+^ transient in the nucleus (Fig. 2B,D and Table S3); this is consistent with the predicted role played by the nuclear envelope - in continuity with the ER lumen - as a major source for nuclear and perinuclear Ca^2+^ spiking (Capoen *et al*., 2011).

The second drug we used was VAC1, an inhibitor of SNARE-dependent vesicle fusions. VAC1 treatment is known to reduce cargo and membrane delivery to the vacuole, leading to an overall reduction in the vacuole size (Dünser *et al*., 2022), thereby potentially affecting the Ca^2+^ homeostasis. After pre-treatment of *L. japonicus* roots with VAC1, the phase 1 Ca^2+^ peak in response to CO4 was strongly reinforced (Fig. 2A,C and Supplementary Fig.S2), suggesting that VAC1-dependent alterations in vacuolar function reduced Ca^2+^ uptake from the cytosol. Nevertheless, the nuclear Ca^2+^ transient was largely unaffected in VAC1 pre-treated roots (Fig. 2B,D and Table S3), indicating that the vacuole has a minor (if any) role in the generation of symbiotic Ca^2+^ signals in the nucleus.

In short, our pharmacological analyses confirmed on the one hand the crucial role of the ER in the generation of nuclear Ca^2+^ signals in response to short-chain chitin oligomers (CO4). On the other hand, our data also demonstrated that the early and strong Ca^2+^ peak triggered by CO4 in the cytosol results from the combined action of a Ca^2+^ influx from the apoplast and sequestration in the vacuole.

### Nuclear and cytosolic Ca^2+^ signatures triggered by chitin-derived oligomers differentially depend on the Common Symbiotic Signalling Pathway

To evaluate the role of the CSSP in triggering the observed Ca^2+^ signals in *L. japonicus* roots upon perception of chitin-derived molecules, we compared our analyses in wild-type *L. japonicus* and mutants for *LjSYMRK* and *LjCASTOR*, two well-characterised components of the CSSP (Oldroyd, 2013; Choi *et al*., 2018; Radhakrishnan *et al*., 2020).

Phase 1 of the cytosolic Ca^2+^ transient was found to be maintained in the CSSP mutants, but with slight differences among stimuli and genotypes. Firstly, the CO4-induced cytosolic Ca^2+^ peak was significantly reduced in *castor* compared to both the wild-type and *symrk* (Supplementary Fig. S2A-B, Dataset S3 and Table S3). Secondly, the CO8-induced cytosolic Ca^2+^ peak showed a limited increase in both *symrk* and *castor*, compared to the wild-type (Supplementary Fig. S2E-F and Table S3). Lastly, no differences could be identified in the mycLCO-induced cytosolic Ca^2+^ peak in phase 1, which was anyway rather limited also in the wild-type (Supplementary Fig. S2I-J and Table S3). By contrast, a slight reduction was detected in the phase 2 Ca^2+^ elevation in response to both CO4 and mycLCOs in both the CSSP mutants in comparison with the wild-type, whereas the Ca^2+^ traces measured in response to CO8 were nearly superimposable (Supplementary Fig. S2 and Table S3).

The differences between genetic backgrounds became more apparent when focusing on nuclear Ca^2+^ transients (Fig. 3). Indeed, the phase 2 broad dome-shaped Ca^2+^ elevation was absent in CO4- and mycLCO-treated mutants, but only weakly reduced upon CO8 application. This is further supported by the quantification of mobilised Ca^2+^ in terms of integral [Ca^2+^] and time duration of the response above an arbitrary [Ca^2+^] threshold (width) (Fig. 3A-D, I-L and Table S3). Conversely, the initial and steep increase in nuclear [Ca^2+^] triggered by the three stimuli was retained in *symrk* and *castor* mutants. Together, these results suggest that a functional CSSP is not required for the cytosolic Ca^2+^ influx of phase 1, even if it appears to modulate it. The absence or reduction of the phase 2 nuclear and cytosolic Ca^2+^ elevation, respectively, in CSSP mutants were particularly intriguing because they mirrored the lack of nuclear and perinuclear Ca^2+^ spiking described in literature for mutants of the orthologous genes in *M. truncatula* (Genre *et al*., 2013; Feng *et al*.,2019). We therefore hypothesised that the dome-shaped Ca^2+^ elevation recorded in phase 2 by our aequorin-based analyses of Ca^2+^concentration in the whole root system may correspond to the sum of individual and non-synchronous Ca^2+^-spiking events triggered in single root cells and previously described in studies based on fluorescent GECIs (Genre *et al*., 2013; Kelner *et al*., 2018; Feng *et al*.,2019). Furthermore, by combining these observations with the results of our EGTA treatments, we concluded that the nuclear Ca^2+^ signalling triggered by CO4 and mycLCOs is composed of a CSSP-independent Ca^2+^ influx from the apoplast during phase 1 and a CSSP-dependent, nuclear-envelope generated Ca^2+^ elevation during phase 2, that we speculated to correspond to nuclear Ca^2+^-spiking.

**Fig. 3.**
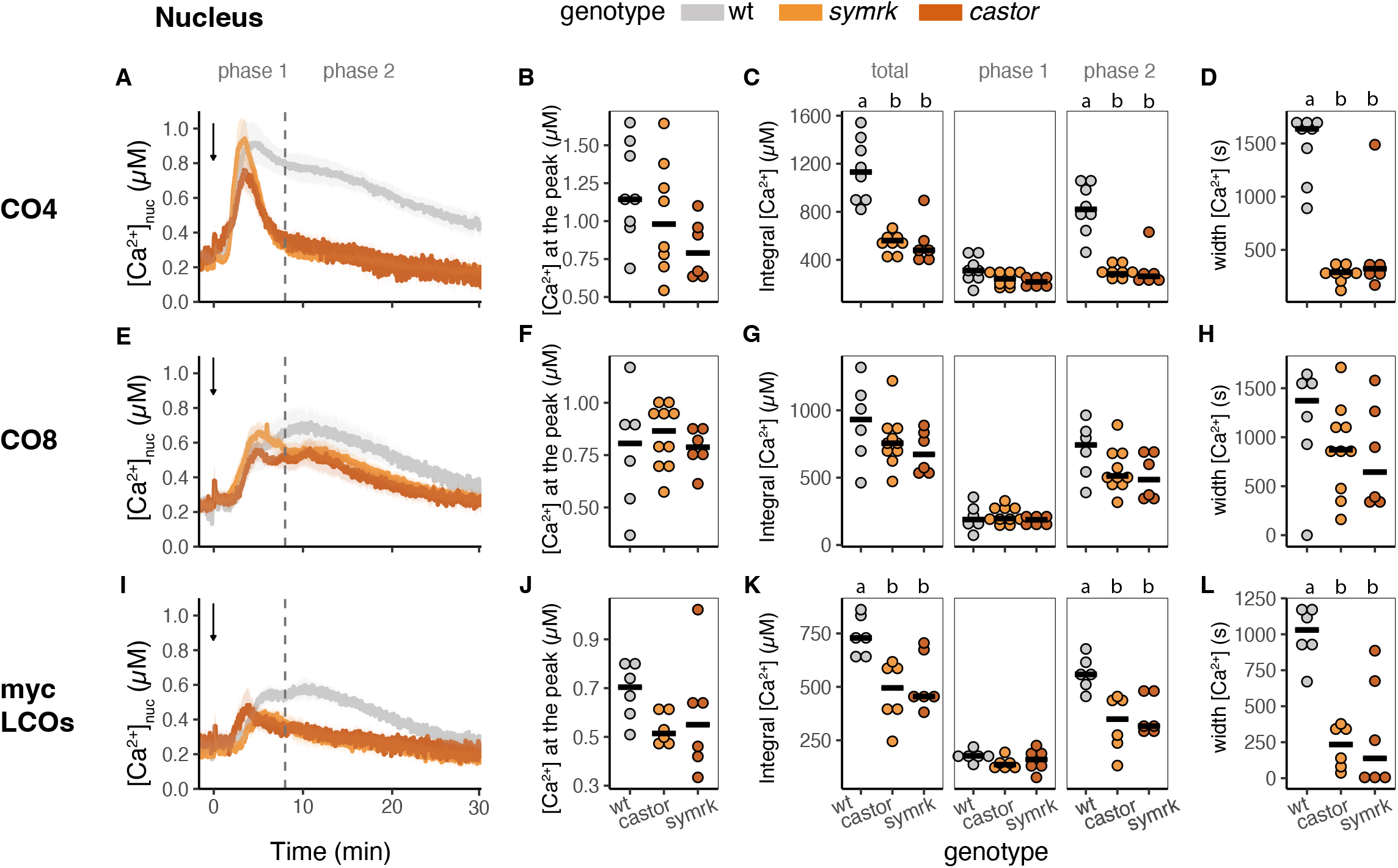
Monitoring of free nuclear [Ca^2+^] in 5 mm-long root segments from composite plants of *L. japonicus* Gifu wt (grey), *castor* (dark orange), *symrk* (light orange). Ca^2+^ measurements were conducted in response to 10^−7^ M CO4 (A-D), 10^−6^ M CO8 (E-H), 10^−7^ M mycLCOs (I-L). In A, E, I, data are presented as means ± SE (shading) of n≥6 traces from at least 3 different composite plants (independent transformation). Arrows indicate the time of stimulus injection (time 0). The dashed line separates phase 1 (0-8 minutes after stimulus) and phase 2 (8-30 minutes after stimulus). In B, F, J, dots represent the maximum [Ca^2+^] for each trace in the whole run. In C, G, K, dots represent the integrated [Ca^2+^] for each trace in the whole run and in the two different phases. In D, H, L, dots represent the Ca^2+^ transient width, in terms of the time interval in which [Ca^2+^] exceeds the arbitrary threshold of 0.4 μM. The black line represents the median of each group. Different letters indicate statistically significant differences among groups, according to Kruskal-Wallis test followed by Dunn’s post-hoc correction (p-value <0.05).

### Cameleon-based analyses of nuclear Ca^2+^ oscillations confirm a multiphasic response to CO4

In order to test this hypothesis, we deployed an alternative set of experiments based on the fluorescent Ca^2+^ probe NupYC2.1, expressed in *Medicago truncatula* root organ cultures. When challenged with 10^−7^ M CO4, root atrichoblasts displayed a cell-autonomous range of Ca^2+^-spiking signals (Fig. 4A,B and Dataset S5), in line with literature data (Genre *et al*., 2013). In more detail, we recorded 30-minute-long traces of at least 9 active atrichoblasts from 8 independent root samples, and the responding cells typically displayed an early elevation in [Ca^2+^] of variable height, shape and duration, but always embraced within the first 8 minutes from treatment. Subsequently, a series of peaks in [Ca^2+^] appeared, also in this case with a very broad variability in terms of peak number, frequency and regularity.

**Fig. 4.**
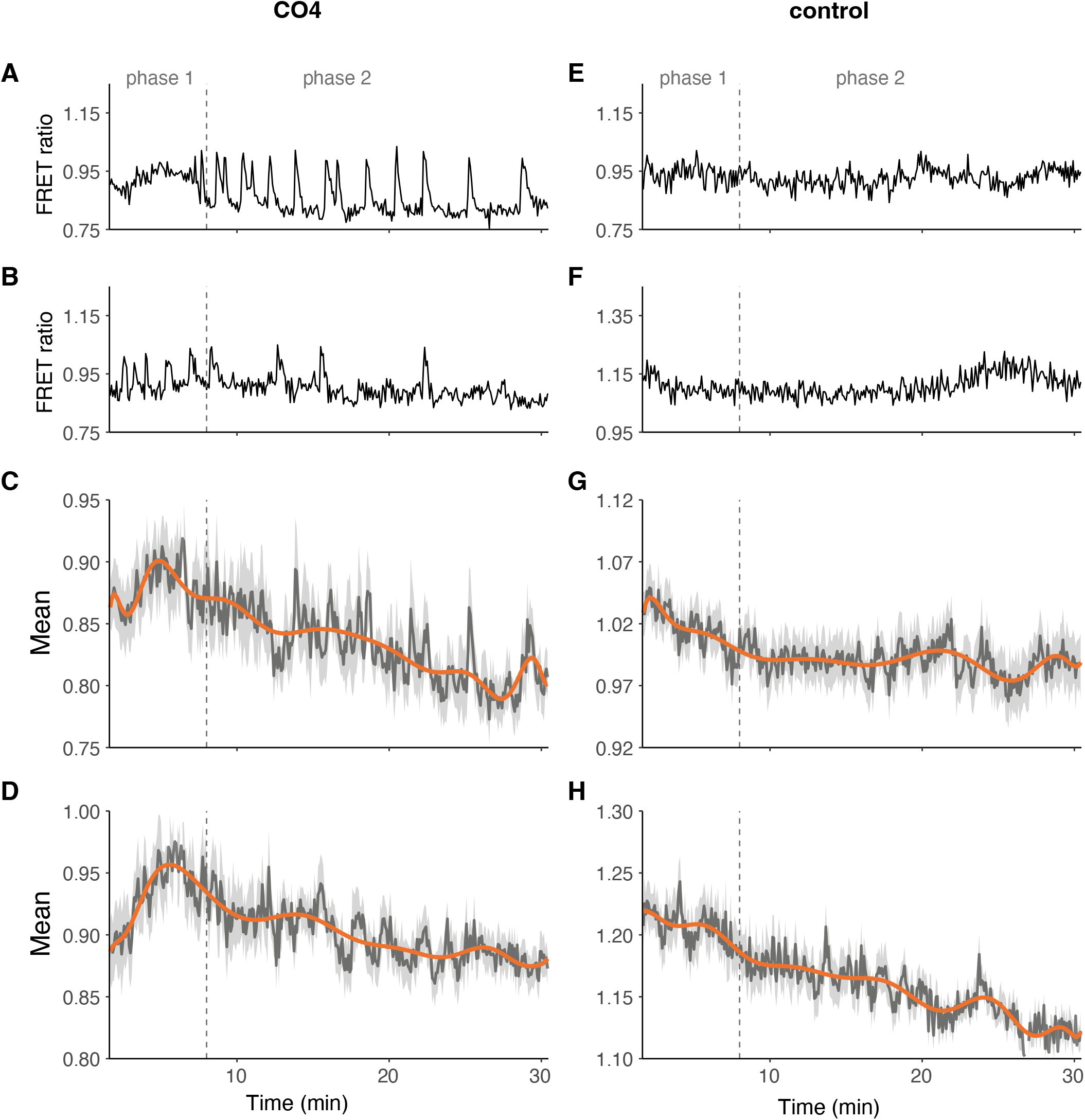
Traces of nuclear Ca^2+^ in root atrichoblasts of *M. truncatula*. A,B,E,F show representative nuclear Ca^2+^ profiles (expressed as YFP/CFP FRET ratio) of individual cells from treated (10^−7^ M CO4) and untreated (control) root segments. CO4 treatment triggered intense oscillations (spiking) in nuclear Ca^2+^ levels. C,D,G,H show polynomial curves (orange) fitting the average values (dark grey) of the FRET traces from the responding cells (light grey) of two independent roots, treated (C,D) or not (G,H) with CO4. The polynomial curves of CO4-treated roots display an initial maximum within the first 8 minutes and a second less pronounced elevation in the following period. A minimum of 9 nuclei were imaged for each root.

In order to compare this cameleon-based, single cell imaging of nuclear Ca^2+^ oscillations with aequorin-based whole-root analyses, we generated average traces combining the signals acquired from all responding cells for each root and a polynomial curve fitting data points of the resulting average trace for each root (Fig. 4C,D and Dataset S1). The resulting Ca^2+^ curve showed an initial elevation during the first 8 minutes after treatment, followed by a second, broader shoulder, with a remarkable resemblance to the traces obtained with our aequorin-based analyses of nuclear [Ca^2+^]. This provided convincing support to our hypothesis that whole-root records by the bioluminescence Ca^2+^ probe correspond to the summation of a population of Ca^2+^ signals from individual cells, as recorded using fluorescence-based Ca^2+^ reporters.

### The initial phase of the CO4- and CO8-induced cytosolic and nuclear Ca^2+^ changes relies on the LysM receptor CERK6

The pharmacological approach and the use of CSSP mutants in the aequorin-based Ca^2+^measurement assays in *L. japonicus* roots challenged with chitin-derived oligomers allowed to abolish either the whole Ca^2+^ response (Fig. 2) or phase 2 only (Fig. 3), but not phase 1 alone. Since phase 1 Ca^2+^ peak is particularly strong upon CO8 and CO4 treatment and a steep cytosolic Ca^2+^ influx is known to play a crucial role in plant immunity (Ranf *et al*., 2012; Monaghan *et al*., 2015; Kutschera *et al*., 2019; Thor *et al*., 2020), we analysed Ca^2+^ responses in the *L. japonicus cerk6* mutant background. CERK6 is a LysM-receptor kinase with high affinity for chitin oligomers and, according to previous reports, *cerk6* plants are unable to mount the immunity response against purified fungal molecules and show increased susceptibility to pathogen infection (Boszoki *et al*., 2017 and 2020). In our hands, CO4 and CO8-induced cytosolic and nuclear Ca^2+^ responses were partially impaired in *cerk6* mutants compared to wild-type and *symrk* mutants (Fig. 5 and Supplementary Dataset S4). Even if only about 30% of *cerk6* root fragments displayed Ca^2+^ signals in response to our treatments (compared to a responsiveness close to 100% in the other genetic backgrounds), all responding *cerk6* plants showed a very similar Ca^2+^ transient, whose timing was consistent with the maintenance of phase 2 only, as recorded in wild-type plants. This observation provided important additional insight into the biphasic nature of the Ca^2+^ responses that we recorded in the wild-type. We can in fact conclude that these include a CERK6-dependent, CSSP-independent phase 1, embracing a major Ca^2+^ influx from the apoplast, and a CERK6-independent, CSSP-dependent phase 2, corresponding to nuclear/perinuclear Ca^2+^-spiking.

**Fig. 5.**
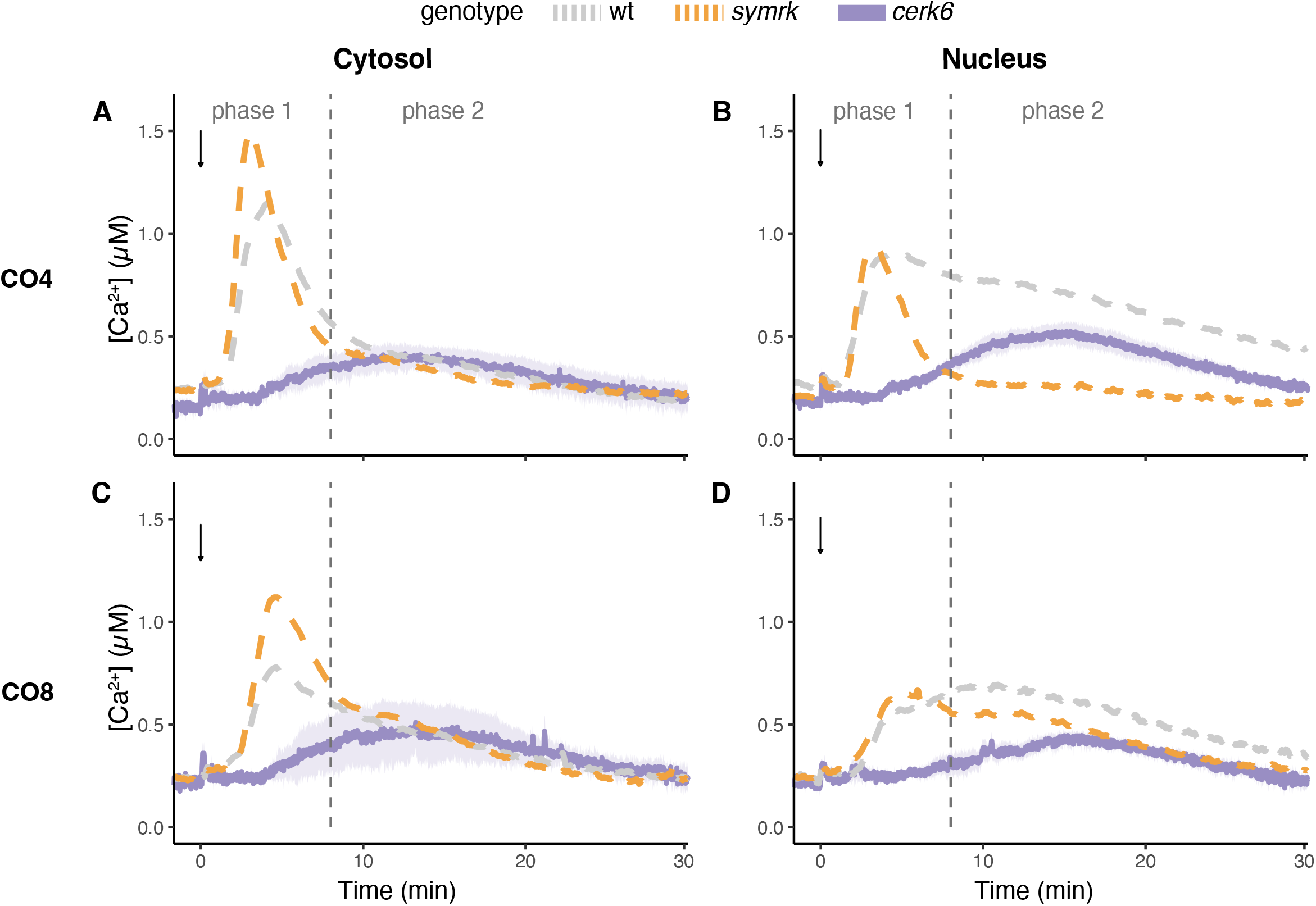
Monitoring of free cytosolic and nuclear [Ca^2+^] in 5 mm-long root segments from composite plants of *L. japonicus cerk6* (purple). Ca^2+^ measurements were conducted in the cytosol (A,C) and nucleus (B,D) in response to 10^−7^ M CO4 (A,C), 10^−6^ M CO8 (B,D). Data are presented as means ± SE (shading) of n≥3 traces from at least 3 responsive composite plants (independent transformation). The Ca^2+^ responses measured in the Gifu wt (grey) and *symrk* (light orange) are shown as moving average (dashed line) for comparison. Arrows indicate the time of stimulus injection (time 0). The dashed line separates phase 1 (0-8 minutes after stimulus) and phase 2 (8-30 minutes after stimulus).

### Fungal elicitor concentration affects the amplitude of the phase 1 Ca^2+^ elevation and the activation of immunity marker genes

It has previously been demonstrated that intracellular Ca^2+^ changes in response to rhizobial Nod-factors are concentration-dependent (Shaw et al, 2003). To test if this was also the case for fungal chitin oligomers, we monitored cytosolic and nuclear Ca^2+^ signals in *L. japonicus* roots challenged with serial dilutions of CO4 and CO8 (Fig. 6, Supplementary Fig. S3 and Dataset S6). Intriguingly, we found that the amplitude of the phase 1 peak in [Ca^2+^] is strongly dependent on the concentration for both CO4 and CO8. Conversely, phase 2 was not apparently affected by the working dilutions. In more detail, 10^−6^ M and 10^−7^ M CO4 generated comparable Ca^2+^ signals, suggesting that the system is already saturated at 10^−7^ M. By contrast, the phase 1 Ca^2+^ peak triggered by 10^−8^ M CO4 was reduced in the cytosol, but unchanged in the nucleus. Moreover, 10^−9^ M CO4 did not trigger any response in the cytosol, while the nuclear phase 2 elevation persisted (Fig. 6A-B). In line with these observations, the treatment of NupYC2.1 *M. truncatula* ROCs with 10^−9^ M CO4 resulted in nuclear Ca^2+^ traces that often lacked the phase 1 Ca^2+^ influx (Supplementary Fig. S4 and Dataset S7) but retained phase 2 Ca^2+^ spiking, albeit overall less pronounced than upon 10^−7^ M CO4 treatments (Fig. 4).

**Fig. 6.**
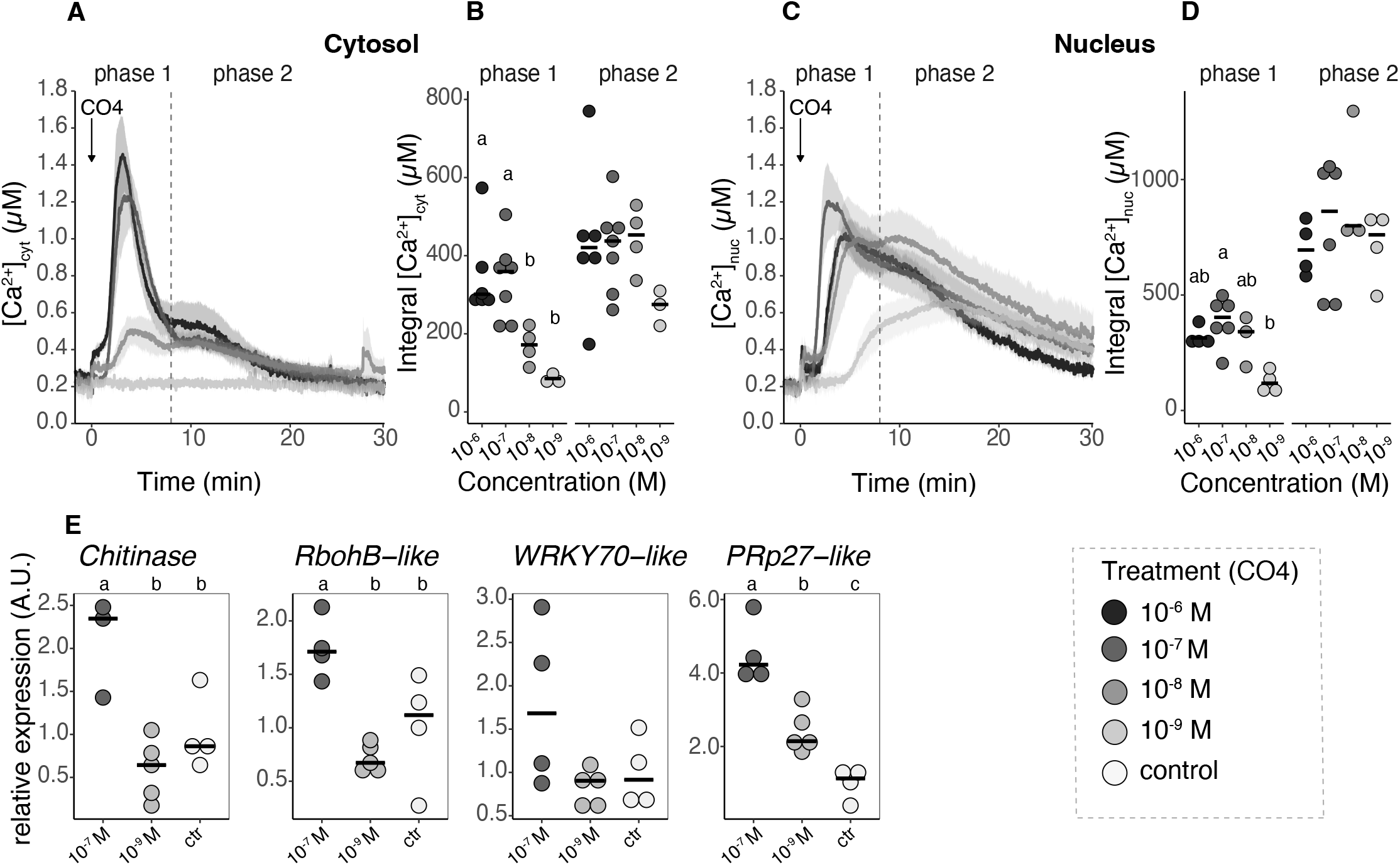
The effect of serial dilutions of CO4 on the induced intracellular Ca^2+^ changes (A-D) and on the activation of immunity marker genes (E). In A, changes in free cytosolic and nuclear [Ca^2+^] were measured in 5 mm-long root segments from composite plants of *L. japonicus* in Gifu wt in response to progressive dilutions of CO4 (10^−6^ M black trace, 10^−7^ M dark grey trace, 10^−8^ M grey trace, 10^−9^ M light grey trace). In A and C, data are presented as means ± SE (shading) of n≥3 traces from at least 3 different composite plants (independent transformation). Arrows indicate the time of stimulus injection (time 0). The dashed line separates phase 1 (0-8 minutes after stimulus) and phase 2 (8-30 minutes after stimulus). In B and C, dots represent the integrated [Ca^2+^] for each trace in the two different phases. The black line represents the median of each group. In E, gene expression analysis by qRT-PCR of *LjChitinase* (Lj5g3v1961260), *LjRbohB-like* (Lj6g3v1549190), *LjWRKY70-like* (Lj1g3v1134110), *LjPRp27-like* (Lj5g3v2112200) relative to the reference gene *LjUbiquitin. L. japonicus Gifu* seedlings were treated for 1 h with solvent control (white), 10^−9^ M CO4 (light grey) or 10^−7^ M CO4 (dark grey) solutions. For each gene, expression is normalized to the control group average. Each dot represents a biological replicate, which is a pool of roots from 12 different plants. The black line represents the median. Different letters indicate statistically significant differences among groups according to Kruskal Wallis and Dunn’s post hoc tests (B, D) or to ANOVA test followed by Tukey’s post-hoc correction (E) (p-value<0.05).

To investigate the physiological relevance of phase 1 Ca^2+^ peak and its possible link with plant defence system, we decided to test to what extent 10^−7^ M and 10^−9^ M CO4 trigger the expression of the immunity marker genes *LjChitinase, LjRbohB-like, LjWRKY70-like, LjPRp27-like* (Bozsoki *et al*., 2017). Indeed, our highest concentration of CO4 triggered the expression of all four immunity markers as soon as 1 h after treatment. By contrast, 10^−9^ M CO4 failed to activate the above-mentioned genes, with the only exception of a slight but significant induction of *PRp27-like* (Fig. 6E and Supplementary Dataset S8). Intriguingly, according to LotusBase ExpressionAtlas (Mun *et al*., 2016; Kamal *et al*., 2020), *LjPRp27-like* shows a peculiar expression pattern with strong and specific gene activation in both Ralstonia-infected plants and AM-colonised roots (Supplementary Fig. S5). This expression profile is unique among the tested immunity marker genes (Lotus Base) and could provide an explanation for the activation of *LjPRp27-like* gene by the low concentration of CO4.

Altogether, these results are consistent with our analysis of Ca^2+^ signals and indicate a dose-dependent regulation of both Ca^2+^-mediated signalling and immunity-related gene expression.

## DISCUSSION

In this work, we identified complex and multiphasic compartment-specific Ca^2+^ signatures activated in response to different fungal signals in *L. japonicus* roots. Aequorin-based Ca^2+^ measurement assays demonstrated that short-chain (CO4), long-chain (CO8) and lipidated (mycLCOs) chitooligosaccharides are all able to induce both cytosolic and nuclear Ca^2+^ transients, that we were able to dissect in two different temporal phases underpinning diverging signalling pathways.

### Chitin oligomers trigger a rapid Ca^2+^ elevation (phase 1) dependent on their concentration and CERK6 activity

We found that all three tested fungal elicitors (CO4, CO8, mycLCOs) are able to induce an early and steep Ca^2+^ increase (phase 1) in both the cytosol and nucleus, independent of the CSSP and the CPA pre-treatment. These data confirm and extend previous reports in soybean and *L. japonicus* cell suspension cultures in response to germinating spore exudates (Navazio *et al*., 2007; Francia *et al*., 2011; Moscatiello *et al*., 2018). The temporal dynamics and shapes of cytosolic phase 1 resemble the well-known Ca^2+^ influx activated by PAMPs that is crucial for mounting plant immunity (Ranf *et al., 2013*, Zipfel and Oldroyd, 2017). We also characterised an analogous phase 1 in nuclear Ca^2+^ signals (and confirmed it at the single-cell level in *M. truncatula* root organ cultures), which appears to be associated with the cytosolic one, suggesting that the same signalling process may extend to the two cell compartments via diffusion of cytosolic Ca^2+^ into the nucleoplasm. The abolishment of the CO4-induced Ca^2+^ elevation in cytosolic phase 1 by EGTA and LaCl3 confirms that a Ca^2+^ influx from the apoplast is at the origin of this process. Intriguingly, the use of VAC1, a recently developed drug affecting membrane delivery to the vacuole and thereby reducing vacuole size (Dünser *et al*., 2022), hinted at the involvement of this extensive compartment in the dissipation of the Ca^2+^ rise in phase 1. The application of the newly designed aequorin-based (Cortese *et al*., 2022) and GCaMP-based (Luo *et al*., 2020; Resentini *et al*., 2021) probes targeted to the plant ER, together with yet-to-develop GECIs for Ca^2+^ measurement/imaging inside the vacuolar lumen, will help uncover the contribution of these Ca^2+^ pools to the generation and dissipation of this phase 1 Ca^2+^ response.

Our analyses of mutant backgrounds indicating the CERK6 dependence, but CSSP independence, of phase 1 signalling in response to chitin oligomers strongly suggests that the phase 1 Ca^2+^ influx is linked to a signal transduction pathway different from the CSSP, in line with previous indications that fungal elicitors may activate multiple parallel signalling cascades (Bonfante and Requena, 2011). Moreover, we observed phase 2 responses in absence of phase 1 under very low concentrations of the fungal elicitors or in the *cerk6* mutants. Similarly, Nod-factor-induced Ca^2+^ influx in *M. truncatula* root hairs has previously been shown to be independent of the CSSP but dependent on the Nod-factor dose (Shaw and Long, 2003) and chemical structure (Morieri *et al*.,2013). Here, CERK6-dependence suggests that phase 1 Ca^2+^ signals are associated with plant immunity (Bozsoki *et al*., 2017), and this is further supported by the activation of immunity marker genes by 10^−7^ M (but not 10^−9^ M) CO4. However, we cannot exclude a role for phase 1 Ca^2+^ influx also in symbiotic signalling. Indeed, the absence of a mycorrhizal phenotype of *cerk6* was previously observed at a single time point (Bozsoki *et al*., 2017), while a deeper analysis of AM colonisation in *MtLyk9* mutants (the putative CERK6 closest homolog in *M. truncatula)* showed a weak reduction compared to wild-type plants (Feng *et al*., 2019; Gibelin-Viala *et al*., 2019). Accordingly, about two-third of the *cerk6* root samples tested in our assays did not respond with a clear activation of phase 2 Ca^2+^ elevation.

### Phase 2 of the Ca^2+^ change induced by chitin oligomers largely depends on the CSSP

By using pharmacological pre-treatments and different plant genetic backgrounds, we showed that the prolonged, phase 2-associated, Ca^2+^ elevations share several features with the Ca^2+^ spiking events induced by chitin oligomers and recorded with fluorescent GECIs (Genre *et al*., 2013; Kelner *et al*., 2018; Feng *et al*., 2019). Such Ca^2+^ spiking events are acknowledged to depend on the CSSP and originate from the nuclear envelope, via the concerted activity of Ca^2+^-permeable channels and Ca^2+^ pumps (Zipfel and Oldroyd, 2017; Charpentier, 2018). Here, we showed that *symrk* and *castor* mutants lack this nuclear Ca^2+^ response (phase 2), which is also abolished by the ER-type Ca^2+^-ATPase inhibitor CPA, as previously shown for Nod factor-induced Ca^2+^ spiking (Capoen *et al*., 2011).

We now hypothesise that perinuclear Ca^2+^ spiking, which is known to be simultaneous with nuclear Ca^2+^ spiking (Ehrhardt *et al*., 1996; Kelner *et al*., 2018), is represented in our aequorin-based analyses by the cytosolic phase 2 elevation. The low amplitude of this cytosolic Ca^2+^ transient may be due to the fact that the aequorin chimera (CPK17_G2A_-NES-YA) used in this work measures [Ca^2+^] changes in the bulk cytosol, rather than at microdomains close to the nuclear envelope (Mehlmer *et al*., 2012; Ottolini *et al*., 2014).

Despite being widely considered as canonical fungal PAMPs (Cao *et al*., 2014; Bjornson *et al*.,2021), CO8 have recently been suggested to also act as symbiotic molecules and activate both nuclear Ca^2+^ spiking and the expression of symbiotic genes (Feng *et al*., 2019, Zhang *et al*., 2021). In our experimental setup, CO8 activated a nuclear phase 2 Ca^2+^ response in the wild-type background, apparently similar to the CO4- and mycLCOs-induced ones. However, this response was only weakly dependent on SYMRK and CASTOR, highlighting that CO8-induced Ca^2+^signalling also underlies transduction pathways unrelated to symbiosis.

### Complementary approaches to measure and image intracellular Ca^2+^ can help disentangle the intertwined Ca^2+^ signalling events during plant root-fungus interaction

By comparing data coming from *L. japonicus* roots expressing nuclear aequorin and nuclear-targeted cameleon in *M. truncatula* root organ cultures, we depicted a precise correspondence between the two datasets, eventually leading to significant insights into plant symbiotic signalling. On the one hand, we confirmed the occurrence of a rapid Ca^2+^ change in response to CO4 perception during phase 1. This initial Ca^2+^ elevation induced by short-chain chitin oligomers is recognizable in published Ca^2+^ traces obtained with cameleon-based techniques but has been largely overlooked, most likely due to its quick and irregular appearance, and analogous Ca^2+^ elevations in response to Nod-factor perception have only been investigated in a couple of studies (Shaw *et al*.,2003; Morieri *et al*., 2013). In this frame, our results bring this elusive element of Ca^2+^-mediated symbiotic signalling back in the spotlight, hopefully fostering new research. Our present results show that a combination of cameleon-based Ca^2+^ traces from individual root epidermal cells does mimic the overall dynamics of this initial CO4-induced Ca^2+^ transient as recorded using aequorin, but cannot fully reproduce its kinetic parameters, suggesting that additional cell layers (*e.g*. belonging to the root cortex), may contribute to the global root Ca^2+^ response to chitooligosaccharides. On the other hand, the combination of two investigation methods provided convincing support to our interpretation of the phase 2 elevation in Ca^2+^ concentration as the summation of individual nuclear Ca^2+^-spiking signals. This is a significant advance in our understanding of symbiotic signalling. In fact, the apparent similarity between aequorin-based and cameleon-based averaged traces of Ca^2+^ signals allows a direct comparison between the results obtained with these alternative and very common approaches. This possibility paves the way for high-throughput genetic screenings and will hopefully lead to fruitful synergies and novel intuitions on the way to disentangling the complexity of Ca^2+^-mediated symbiotic signalling in plants.

In conclusion, recent literature has suggested a cross-talk between immunity and symbiosis signalling pathways upon plant perception of root endosymbionts. In this scenario, the prevalence of either pathway depends on a combination of elicitors, receptor competition and cross-reactions among players of the activated signalling cascades (Feng *et al*., 2019, Zhang *et al*., 2021, Feng *et al*., 2021). Here, we suggest that the boundaries between *bona-fide* pathogenic and symbiotic fungal signals are less clear-cut than previously thought, since all the tested molecules could activate parallel pathways that converge in a multi-phasic Ca^2+^ signal. In fact, we could dissect independent components of the Ca^2+^ responses based on the genetic background, time phase and elicitor concentration. Moreover, we correlated the activation of immunity marker genes with the presence of the phase 1 Ca^2+^ influx, via its critical dependence on the concentration of the elicitor, suggesting that the concentration of chitin-derived molecules plays a crucial role in the activation of different plant responses to root-interacting microbes. Future research will be needed to further characterise the potential role of this Ca^2+^ influx in AM symbiosis.

## Supplementary Data

Fig. S1. Genetic map and intracellular localization of the cytosolic and nuclear YFP-aequorin chimeras in *L. japonicus* roots.

Fig. S2. CO4-, CO8- and mycLCOs-induced cytosolic Ca^2+^ responses in *L. japonicus* CSSP mutant backgrounds.

Fig. S3. Cytosolic and nuclear Ca^2+^ responses to serial dilutions of CO8 in *L. japonicus* roots.

Fig. S4. Traces of nuclear Ca^2+^ concentration in root atrichoblasts of *M. truncatula* under 10^−9^ M CO4 treatment.

Fig. S5. Expression profile of the immunity marker genes retrieved from LotusBase ExpAt.

Table S1. List of primers used in this study.

Table S2. List of plasmids used in this study.

Table S3. Extended outcomes of the statistical analysis for all experiments in this manuscript. Dataset S1. Script to reproduce all statistical analysis and figures in RStudio environment. Dataset S2. Raw data for Fig. 1.

Dataset S3. Raw data for Fig. 2.

Dataset S4. Raw data for Fig. 3, Fig. 5 and Supplementary Fig. S5.

Dataset S5. Raw data for Fig. 4.

Dataset S6. Raw data for Fig. 6A-D and Supplementary Fig. S3.

Dataset S7. Raw data for Supplementary Fig. S4.

Dataset S8. Raw data for Fig. 6E.

## Acknowledgements

We thank Ute Vothknecht (University of Bonn, Germany) for the CPK17_G2A_-NES-YA and NLS-YA plasmids, Marco Incarbone and Mattia Donà (Gregor Mendel Institute, Vienna, Austria) for the destination vector, and Sébastien Fort (CNRS, Paris, France) for the mycLCOs. We are grateful to Lene Heegaard Madsen, Simona Radutoiu and Jens Stougaard (Aarhus University, Denmark) for kindly providing *L. japonicus symrk, castor* and *cerk6-1* mutants, and to Myriam Charpentier (John Innes Centre, Norwich, UK) for fruitful discussions. The technical assistance of the Imaging Facility and the Plant Genome Editing Facility of the Department of Biology (University of Padova, Italy) is gratefully acknowledged.

## Author contributions

LN and MG conceived the study and designed research; FB and MG conducted molecular cloning; FB performed intracellular localization studies and gene expression analysis; FB and EO performed aequorin-based Ca^2+^ measurements; AC and FB performed cameleon-based Ca^2+^ assays; MG and FB analysed data, conducted statistical analyses and data visualisation; AC, IS and FB conducted data analysis and modelling of Ca^2+^ imaging data; FB, MG and LN wrote the article; AG designed some experiments and contributed to the discussion and editing of the article; JKV provided materials and contributed to the discussion. All authors read and approved the final manuscript.

## Data availability

The datasets used in this study are available in the Supplementary data.

