## Supplementary material for "Unlocking the multiphasic nature of intracellular calcium signatures triggered by fungal signals in *Lotus japonicus* roots": Dataset S1: Dataset_S1.Rmd

````
---
title: "Dataset_S1"
output:
  html_document: default
  pdf_document: default
date: "2023-01-18"
---

###Figure 1

##Loading and tidying aequorin files_Figure 1
```{r}
##Before starting, move both the .csv files and this R Markdown file in the same folder

##check the working directory
getwd()

##install the following packages and/or load them
library(ggExtra)
library(tidyverse)
library(multcomp)
library(ggpubr)
library(rstatix)
library(patchwork)
library(writexl)

##path to the supplementary file "Dataset_S2_raw_data_Fig1"
data <- read.csv(file= "Dataset_S2_raw_data_Fig1.csv", header=T,sep = ";", na.string=c("na", "NA", "null")) 

str(data)
data_tidy <- pivot_longer(data, cols=2:ncol(data), names_to= "condition", values_to = "calcium")

data_tidy <- data_tidy %>% separate(condition, sep="_", c('compartment', 'treatment', 'genotype', 'replicate'))
data_tidy$compartment <- as.factor(data_tidy$compartment)
data_tidy$treatment <- as.factor(data_tidy$treatment)
data_tidy$genotype <- as.factor(data_tidy$genotype)
data_tidy$replicate <- as.factor(data_tidy$replicate)


data_tidy$treatment <- recode_factor(data_tidy$treatment, co4="CO4", co8="CO8", myclco="mycLCO")

data_tidy$compartment <- recode_factor(data_tidy$compartment, cya="cytosol", nls="nucleus")

data_tidy <- na.omit(data_tidy)
head(data_tidy)

data_tidy <- data_tidy %>% 
  mutate (time_after_treatment=time_points-1.667)

```

##Data frame with stats: mean, standard deviation, standard error_Figure 1
```{r}
data_stats  <- data_tidy %>% 
  group_by(time_after_treatment, genotype, compartment, treatment) %>% 
 mutate(mean = mean(calcium), 
            stdev = sd(calcium),
            sterr = sd(calcium)/sqrt(length(calcium)),
   UI = paste(compartment, treatment, genotype, replicate, sep = '_')) 
```

##Traces per each compartment_Figure 1
```{r}
data_stats%>% 
  filter(genotype=="wt") %>%
  split(.$compartment) %>% 
    map(
~ggplot (., aes(time_after_treatment, mean,   color=treatment, fill=treatment)) +
  geom_line(lwd=1) +
  scale_colour_manual(values=c("#1f78b4", "#b2df8a", "#a6cee3", "#cccccc")) + 
  scale_fill_manual(values=c("#1f78b4", "#b2df8a", "#a6cee3", "#cccccc")) +
  geom_ribbon(aes(ymin=mean-sterr, ymax=mean+sterr),alpha=0.2,   colour = NA) +  
  scale_y_continuous(breaks=seq(0, 1.5, 0.2), expand = c(0,0), limits= c(0,1.5))+
  theme_classic() +
  annotate("segment", x = 0, y = 0.9, xend = 0, yend = 0.8,lineend = "round", linejoin = "round",
         arrow = arrow(type = "closed", length = unit(0.02, "npc"))) +
  annotate ("text", x=20, y=0.93, label=paste0(unique(.$compartment)), colour="black")+
  labs(y=paste0(unique(.$compartment)," Ca2+ concentration"), x="Time (min)") +
    theme(legend.position="none") 
) -> trace_wt_compartment

trace_wt_compartment[[1]]+
  trace_wt_compartment[[2]]+
      plot_annotation(tag_levels = 'A') +  
  plot_layout(guides = 'collect', ncol = 2) -> plot_fig_compartment
```

##Scatterplot per each compartment_Figure 1
```{r}
data_max_peak  <- data_stats %>% 
  filter (time_after_treatment > 1) %>%  ##this parameter is to avoid 
  ##to consider a peak when is due to some "touch" event
  filter (time_after_treatment <20) %>%  
   group_by(genotype, compartment,  treatment, replicate) %>% 
  summarise(max_peak = max(calcium)) %>% 
      mutate(UI = paste(genotype,compartment, treatment, replicate, max_peak, sep = '_'))

data_tidy <- data_tidy %>% 
 mutate(UI = paste(genotype, compartment, treatment, replicate, calcium, sep = '_'))

data_merged <- dplyr::left_join(data_max_peak, data_tidy, 
                                by = c("UI", "genotype", "compartment", "treatment", "replicate")) 
## new dataframe with time and maxpeak intensity
##scatterplot max calcium and time - P3 P4 (panel C and D)

data_merged %>%  
  filter(treatment !="ctr") %>% 
  split(.$compartment)  %>% 
  map(
  ~ggplot(., aes(x=time_after_treatment, y=max_peak, fill=treatment, colour=treatment))+
            scale_colour_manual(values=c("#1f78b4", "#b2df8a", "#a6cee3")) +
  scale_fill_manual(values=c("#1f78b4", "#b2df8a", "#a6cee3")) +
   geom_point(size=3) +
      scale_x_continuous( limits= c(2.5,17))+
    scale_y_continuous (limits= c(0.4, 2.0, 0.5)) +
              theme_classic()    +
  labs(y= Ca^"2+"~peak~(mu*M), x= "Time (min)") +
      theme(#axis.title.x=element_blank(), 
            legend.position = "none",
            axis.text=element_text(size=11),
            panel.border = element_blank(),
            panel.grid.minor = element_blank(),
        strip.background = element_blank()) 
 )-> list_scatter
  
P3 <- ggMarginal(list_scatter[[1]], groupColour = TRUE, groupFill = TRUE) 
P4 <- ggMarginal(list_scatter[[2]], groupColour = TRUE, groupFill = TRUE) 

###----Figure 1-----#####
trace_wt_compartment[[1]] + 
trace_wt_compartment[[2]] +
  P3+
  P4+
  plot_annotation(tag_levels = 'A') +  
  plot_layout(guides = "auto", ncol = 2) #-> plot_fig1_scatterplot
```

##check normality and homogeneity of variance on max_peak_Figure 1
```{r}

# Build the linear model
lm(max_peak ~ treatment, data = data_max_peak) -> model 

 # Create a QQ plot of residuals and check for normality
ggqqplot(residuals(model))
shapiro_test(residuals(model))

##check Homogeneity of variance 
plot(model, 1)
model  %>% levene_test(max_peak ~ treatment)
##there is no homogeneity of variance, then we will apply a non-parametric test

##there is no homogeneity of variance, then we will apply a non-parametric test

## Kruskal Wallis on max peak
data_max_peak %>% 
  group_by(compartment) %>% 
  kruskal_test(max_peak~ treatment ) #%>% ####remove the hashtags if you want to create an excel file with the stats summary
 #write_xlsx(., path = " Krusk_max_peak_1.xlsx")

## Dunn's post-hoc test
data_max_peak %>% 
   group_by(compartment) %>% 
  dunn_test(max_peak~ treatment , p.adjust.method = "fdr")  #%>% ####remove the hashtags if you want to create an excel file with the stats summary
 #write_xlsx(., path = "Dunns_max_peak_1.xlsx")

## Box plot on maximum peak 
ggplot(data_max_peak, aes(x=treatment, y=max_peak, color=treatment)) + 
  scale_fill_manual(values=c("#808080", "#756bb1","#bdd7e7", "#6baed6", "#3182bd")) +
    geom_boxplot(position=position_dodge(0.9), outlier.shape = NA, lwd=0.4)  +
  theme_classic() +
  theme(axis.title.x = element_text(size=14, face="bold"),
axis.title.y = element_text(size=14, face="bold"),
axis.text.y =element_text(size=12), axis.text.x =element_text(size=12)) +
  scale_color_manual(values=c("#808080", "#756bb1","#bdd7e7", "#6baed6", "#3182bd")) +
geom_point(position=position_jitterdodge(dodge.width=0.9), size= 1.8) +
  facet_wrap (~compartment)

```
###Figure 2
##Loading and tidying aequorin files_Figure 2
```{r}
#read the csv file
  data <- read.csv("Dataset_S3_raw_raw_Fig2.csv", header=T,sep = ";", na.string=c("na", "NA", "null"))
str(data)
data_tidy <- pivot_longer(data, cols=2:ncol(data), names_to= "condition", values_to = "calcium")

data_tidy <- data_tidy %>% separate(condition, sep="_", c('compartment', 'treatment', 
                                                          'genotype', 'drug', 'replicate'))

#consider compartment, treatment, genotype, drug, replicate as factors
data_tidy$compartment <- as.factor(data_tidy$compartment)
data_tidy$treatment <- as.factor(data_tidy$treatment)
data_tidy$genotype <- as.factor(data_tidy$genotype)
data_tidy$drug <- as.factor(data_tidy$drug)
data_tidy$replicate <- as.factor(data_tidy$replicate)


data_tidy$treatment <- recode_factor(data_tidy$treatment, co4 = "CO4", co8 = "CO8", 
                                     mycLCO ="mycLCO")

data_tidy$compartment <- recode_factor(data_tidy$compartment, cya="cytosol", nls="nucleus")

data_tidy <- na.omit(data_tidy)
summary(data_tidy)

data_tidy <- data_tidy %>% 
  filter(treatment != "ctrDmso") %>% 
  filter(treatment != "ctrEtoh") %>% 
  droplevels()

data_tidy$drug <- recode_factor(data_tidy$drug , egta = "EGTA",LaCl3 = "LaCl",
                                VAC1 = "VAC1", vac1 ="VAC1")

data_tidy <- data_tidy %>% 
  mutate (time_after_treatment=time-1.667)
```

##database with statistics: mean, standard deviation and standard error_Figure 2
```{r}
data_stats  <- data_tidy %>% 
  group_by(time_after_treatment, genotype, compartment, drug, treatment) %>% 
 mutate(mean = mean(calcium), 
            stdev = sd(calcium),
            sterr = sd(calcium)/sqrt(length(calcium)),
   UI = paste(compartment, treatment, genotype, drug, replicate, sep = '_')) 

data_tidy_with_baseline <-  data_tidy  %>% 
  filter (time_after_treatment < 0) %>% 
    group_by(genotype, compartment, drug, treatment, replicate) %>% 
    mutate(mean_before_treatment =  mean(calcium),
    UI = paste(compartment, treatment, genotype, drug, replicate, sep = '_'))

data_tidy_with_baseline %>% 
ungroup() %>%
dplyr::select(UI, mean_before_treatment) -> selection

#mergeCols <- c("UI")
joined_tables <- dplyr::left_join(data_stats, selection, by = "UI") 
## It creates many row's duplicate
 joined_tables <- joined_tables %>% 
   distinct() %>%  ## removing duplicates
  mutate(delta_calcium = calcium - mean_before_treatment)
```

##aequorin traces drugs - (panel A and B)_Figure 2
```{r}
data_stats$drug <- factor(data_stats$drug, levels = c( "LaCl","EGTA", "none", "CPA", "VAC1"))

data_stats   %>% 
  filter (genotype == "wt") %>% 
   filter (treatment == "CO4") %>% 
    split(.$compartment)  %>% 
  map(
 ~ ggplot (., aes(time_after_treatment, mean,   color=drug, fill=drug)) +
  geom_line(lwd=1) +
 scale_colour_manual(values=c( "#762a83","#af8dc3", "#252525","#7fbf7b", "#1b7837")) +
  scale_fill_manual(values=c( "#762a83","#af8dc3", "#252525","#7fbf7b", "#1b7837")) +
  geom_ribbon(aes(ymin=mean-sterr, ymax=mean+sterr),alpha=0.2, colour = NA) +  
  scale_y_continuous(breaks=seq(0, 2, 0.5), expand = c(0,0), limits= c(0,2))+
  annotate("segment", x = 0, y = 0.9, xend = 0, yend = 0.8,lineend = "round", linejoin = "round",
         arrow = arrow(type = "closed", length = unit(0.02, "npc"))) +
 #ylim (0, 1.25) +
  scale_x_continuous(expand = c(0,0))+
  theme_classic() +
    labs(y= expression(Ca^"2+"~concentration~(mu*M))) +
    theme(axis.title.x=element_blank(),
          legend.position = "none",
          axis.text=element_text(size=11)) 
  ) ->traces_aeq
```

##calculation of delta integral (integral - mean of the integral before adding the stimulus)_Figure 2
```{r}
delta_AUC  <- joined_tables %>% 
  filter (time_after_treatment > 1) %>%   ##this parameter is to avoid to consider a peak when is due to some "touch" event
  group_by(genotype, compartment, drug, treatment, replicate) %>% 
  summarise(delta_integral = sum(delta_calcium)) 

```

##calculation of Max Peak and delta max peak_Figure 2
```{r}
data_max_peak  <- data_tidy %>% 
filter (time_after_treatment > 1) %>%   ##this parameter is to avoid to consider a peak when is due to some "touch" event
    group_by(genotype, compartment, drug, treatment, replicate) %>% 
  summarise(max_peak = max(calcium)) 

##calculation of delta Max Peak
data_delta_max_peak  <- joined_tables %>% 
  filter (time_after_treatment > 1) %>%   ##this parameter is to avoid to consider a peak when is due to some "touch" event
    group_by(genotype, compartment, drug, treatment, replicate) %>% 
  summarise(delta_max_peak = max(delta_calcium)) 
```

##delta AUC (panel D)_Figure 2
```{r}
##This plot is inspired by www.cedricscherer.com/2019/05/17/the-evolution-of-a-ggplot-ep.-1/#polish

delta_AUC$drug <- factor(delta_AUC$drug, levels = c( "LaCl","EGTA", "none", "CPA", "VAC1"))

delta_AUC <- delta_AUC %>% 
  filter (treatment == "CO4") %>% 
  filter (compartment == "nucleus") %>% 
  filter (genotype == "wt") 

delta_AUC$drug <- factor(delta_AUC$drug, levels = c( "LaCl","EGTA", "none", "CPA", "VAC1"))

##mean per each treatment
delta_AUC <- delta_AUC %>% 
  group_by(drug) %>% 
  mutate(mean_delta_integral = mean(delta_integral)) 

g <-  ggplot(delta_AUC, aes(x=drug, y = delta_integral, color = drug)) +
  scale_colour_manual(values=c( "#762a83","#af8dc3", "#252525","#7fbf7b", "#1b7837")) +
               theme_classic() + 
  theme(axis.text=element_text(size=11),
            axis.title.x = element_blank(),
            legend.text = element_text(size = 11)) +
           labs(y= (Delta~Nuclear~integrated~Ca^"2+"~(mu*M))) 
  
##mean of "none" treatment
 wt_delta_integral <-
  delta_AUC %>%
  filter (drug == "none") %>% 
  summarize(avg = mean(delta_integral, na.rm = T)) %>%
  pull(avg)

##adding a segment for the distance between the mean of each treatment and the mean of the "none" treatment
g + 
  geom_segment(
   aes(x=drug, xend = drug,
    y = wt_delta_integral, yend = mean_delta_integral),
   size = 0.7
   ) +
  geom_hline(aes(yintercept = wt_delta_integral), color = "gray70", size = 0.6) +
  geom_jitter(size = 2, alpha = 0.7, width = 0.2) +
  stat_summary(fun = mean, geom = "point", size = 5) -> p4
```
##check normality and homogeneity of variance panel D_Figure 2
```{r}
delta_AUC <- delta_AUC %>% 
 mutate(UI = paste(compartment, drug, treatment, replicate, sep = '_'))

delta_AUC %>% 
  group_by(drug) %>%
  identify_outliers(delta_integral) -> delta_integral_outliers

# Build the linear model, excluding outliers
delta_AUC %>% 
 filter(!UI %in% delta_integral_outliers$UI)  -> delta_AUC

# Build the linear model, excluding outliers
lm(delta_integral ~ drug, data = delta_AUC) -> model 

 # Create a QQ plot of residuals and check for normality
ggqqplot(residuals(model))
shapiro_test(residuals(model))

##check Homogeneity of variance 
plot(model, 1)
model  %>% levene_test(delta_integral ~ drug)

delta_AUC %>% 
   filter (genotype == "wt") %>%
  group_by(compartment) %>% 
  kruskal_test(delta_integral~ drug )  #%>% ####remove the hashtags if you want to create an excel file with the stats summary 
 #write_xlsx(., path = "Krusk_AUC_2D.xlsx")

```
## Dunn's post-hoc test - panel D_Figure 2
```{r}
delta_AUC %>% 
   group_by(compartment) %>% 
  dunn_test(delta_integral ~ drug, p.adjust.method = "fdr")  #%>% ####remove the hashtags if you want to create an excel file with the stats summary 
 #write_xlsx(., path = "Dunns_AUC_2D.xlsx")
```
##plot delta Max Peak cytosol - panel C_Figure 2
```{r}
data_delta_max_peak <- data_delta_max_peak %>% 
  filter (treatment == "CO4") %>% 
  filter (compartment == "cytosol") %>% 
  filter (genotype == "wt") 

data_delta_max_peak$drug <- factor(data_delta_max_peak$drug, levels = c( "LaCl","EGTA", "none", "CPA", "VAC1"))

theme_set(theme_light(base_size = 18, base_family = "Poppins"))

##adding a mean per each genotype
data_delta_max_peak <- data_delta_max_peak %>% 
  group_by(drug) %>% 
  mutate(mean_delta_max_peak = mean(delta_max_peak)) 
          
g <-  ggplot(data_delta_max_peak, aes(x=drug, y = delta_max_peak, color = drug)) +
  scale_colour_manual(values=c( "#762a83","#af8dc3", "#252525","#7fbf7b", "#1b7837")) +
               theme_classic() + 
  theme(axis.text=element_text(size=11),
            axis.title.x = element_blank(),
            legend.text = element_text(size = 11)) +
           labs(y= (Delta~Cytosolic~max~Ca^"2+"~concentration~(mu*M))) 
      
## wt mean
 wt_delta_max_peak <-
  data_delta_max_peak %>%
  filter (drug == "none") %>% 
  summarize(avg = mean(delta_max_peak, na.rm = T)) %>%
  pull(avg)

##distance from wt mean 
g + 
  geom_segment(
   aes(x=drug, xend = drug,
    y = wt_delta_max_peak, yend = mean_delta_max_peak),
   size = 0.7
   ) +
  geom_hline(aes(yintercept = wt_delta_max_peak), color = "gray70", size = 0.6) +
  geom_jitter(size = 2, alpha = 0.7, width = 0.2) +
  stat_summary(fun = mean, geom = "point", size = 5) -> p3

```
##check normality and homogeneity of variance on panel C and Anova test_Figure 2
```{r}
data_delta_max_peak <- data_delta_max_peak %>% 
 mutate(UI = paste(genotype, compartment, drug, replicate, sep = '_'))

data_delta_max_peak %>% 
  group_by(drug) %>%
  identify_outliers(delta_max_peak) -> maxpeak_outliers

# Build the linear model, excluding outliers
data_delta_max_peak %>% 
 filter(!UI %in% maxpeak_outliers$UI)  -> data_delta_max_peak

# Build the linear model, excluding outliers
lm(delta_max_peak ~ drug, data = data_delta_max_peak) -> model 

 # Create a QQ plot of residuals and check for normality
ggqqplot(residuals(model))
shapiro_test(residuals(model))

##check Homogeneity of variance 
plot(model, 1)
model  %>% levene_test(delta_max_peak ~ drug)

# Kruskal wallis test 
data_delta_max_peak %>% 
   filter (genotype == "wt") %>%
  group_by(compartment) %>% 
  kruskal_test(delta_max_peak ~ drug )  #%>% ####remove the hashtags if you want to create an excel file with the stats summary
 #write_xlsx(., path = "Krusk_MaxPeak_2C.xlsx")
```

##Dunn's post-hoc test - 4D_Figure 2
```{r}
data_delta_max_peak %>% 
   group_by(compartment) %>% 
  dunn_test(delta_max_peak ~ drug , p.adjust.method = "fdr")  #%>% ####remove the hashtags if you want to create an excel file with the stats summary
 #write_xlsx(., path = "Dunns_MaxPeak_2C.xlsx")
```

##arrange plots - same y axis_Figure 2
```{r}
###---------Figure 2---------#####
 traces_aeq[[1]] + 
 traces_aeq[[2]] +
 p3+
 p4 +  
   plot_annotation(tag_levels = 'A') +  
  plot_layout(guides = 'collect', ncol = 2) #-> plot_drug_fig2
```

### Figure 3

###Loading and tidying aequorin raw files_Figure 3
```{r}
###your path to the supplementary table "Dataset_S4_raw_data_Fig3-5-S5.csv"
data <- read.csv("Dataset_S4_raw_data_Fig3-5-S5.csv", 
                 header=T,sep = ";", na.string=c("na", "NA", "null")) 
str(data)
data_tidy <- pivot_longer(data, cols=2:ncol(data), names_to= "condition", values_to = "calcium")

data_tidy <- data_tidy %>% separate(condition, c('compartment', 'treatment',                                             'genotype','replicate'))

data_tidy$compartment <- as.factor(data_tidy$compartment)
data_tidy$treatment <- as.factor(data_tidy$treatment)
data_tidy$genotype <- as.factor(data_tidy$genotype)
data_tidy$replicate <- as.factor(data_tidy$replicate)
data_tidy$genotype <- factor (data_tidy$genotype, levels = c("wt", "symrk", "cerk6", "castor"))

data_tidy$treatment <- recode_factor(data_tidy$treatment, co4 = "CO4", co8 = "CO8", myclco="mycLCO", mycLCO="mycLCO")
data_tidy$compartment <- recode_factor(data_tidy$compartment, cya="cytosol", nls="nucleus")

data_tidy <- na.omit(data_tidy) 
head(data_tidy)

data_tidy <- data_tidy %>% 
  mutate (time_after_treatment = time - 1.667)

```

###creating a data frame with stats: mean, standard deviation, standard error_Figure 3
```{r}

data_stats  <- data_tidy %>% 
  group_by(time_after_treatment, genotype, compartment, treatment) %>% 
 mutate(mean = mean(calcium), 
            stdev = sd(calcium),
            sterr = sd(calcium)/sqrt(length(calcium)),
   UI = paste(compartment, treatment, genotype, replicate, sep = '_')) 
```
###plots with traces - wt, symrk, castor_Figure 3
```{r}
data_stats%>%
  filter(genotype!="cerk6")%>%
  filter(compartment=="nucleus") %>% 
  split(.$treatment) %>% 
    map(
~ggplot (., aes(time_after_treatment, mean,   color=genotype, fill=genotype)) +
  geom_line(lwd=1) +
  scale_colour_manual(values=c("#cccccc", "#fe9929",  "#d95f0e")) + 
  scale_fill_manual(values=c("#cccccc", "#fe9929",  "#d95f0e")) +
  geom_ribbon(aes(ymin=mean-sterr, ymax=mean+sterr),alpha=0.2,   colour = NA) +  
  scale_y_continuous(breaks=seq(0, 1.1, 0.2), expand = c(0,0), limits= c(0,1.1))+
  annotate ("text", x=4, y=1, label="Phase 1", colour="#737373")+
  annotate ("text", x=15, y=1, label="Phase 2", colour="#737373")+
  scale_x_continuous(expand = c(0,0))+
  theme_classic() +
    annotate("segment", x = 8, xend = 8, y = 0, yend = Inf, colour="#737373",linetype = "dashed") +
  annotate("segment", x = 0, y = 0.9, xend = 0, yend = 0.8,lineend = "round", linejoin = "round",
         arrow = arrow(type = "closed", length = unit(0.02, "npc"))) +
    annotate ("text", x=1.666, y=0.93, label=paste0(unique(.$treatment)), colour="black")+
  labs(y=paste0(unique(.$compartment)," Ca2+ concentration")) +
    theme(axis.title.x=element_blank()) 
) -> trace_nls

  trace_nls[[1]]+
   trace_nls[[2]]+
   trace_nls[[3]]+
    plot_annotation(tag_levels = 'A') +  
  plot_layout(guides = 'collect', ncol = 3) -> plot_nls_mutsymb

```


###max peak _Figure 3
```{r} 
data_max_peak  <- data_tidy %>% 
  filter (time_after_treatment > 1) %>%  ###this parameter is to avoid 
  ###to consider a peak when is due to some "touch" event
  group_by(genotype, compartment, treatment, replicate) %>% 
    summarise(max_peak = max(calcium)) %>% 
      mutate(UI = paste(compartment, treatment, genotype, replicate, max_peak, sep = '_'))

data_max_peak %>% 
  filter(genotype!="cerk6")%>%
  filter(compartment=="nucleus")%>%
  split(.$treatment)%>%
map(
~ggplot(., aes(x=genotype, y=max_peak, fill=genotype))+
                scale_fill_manual(values=c("#cccccc", "#fe9929",  "#d95f0e")) +
                geom_dotplot(binaxis ="y", stackdir = "center", dotsize = 1.8) + 
                stat_summary(fun = median, fun.min = median, fun.max = median, geom = "crossbar", 
                           width = 0.7) +
                theme_classic()    +
  labs(y= expression(Ca^"2+"~at~the~peak~(mu*M))) +
    annotate ("text", label=paste0(unique(.$treatment)," ",unique(.$compartment)),  
              x=Inf, y=Inf,hjust=1,vjust=1, colour="black")+
  theme(axis.title.x=element_blank(), 
        strip.background = element_blank(),
        panel.border = element_rect(colour = "black", fill = NA))
) -> list_max_all


 trace_nls[[1]]+
   list_max_all[[1]]+
   trace_nls[[2]]+
   list_max_all[[2]]+
   trace_nls[[3]]+
   list_max_all[[3]]+
    plot_annotation(tag_levels = 'A') +  
  plot_layout(guides = 'collect', ncol = 2) -> plot_max_nls_mutsymb
```

###check normality and homogeneity of variance on max_peak_Figure 3
```{r}
data_max_peak%>% 
  filter(genotype!="cerk6") %>% 
  filter(compartment=="nucleus") -> data_max_peak_nls

# Build the linear model
lm(max_peak ~ treatment*genotype, data = data_max_peak_nls) -> model

# Create a QQ plot of residuals and check for normality
ggqqplot(residuals(model))
shapiro_test(residuals(model))

##check Homogeneity of variance 
plot(model, 1)
model  %>% levene_test(max_peak ~ treatment*genotype)
##there is no homogeneity of variance, then we will apply a non-parametric test

# Indentify outliers
data_max_peak_nls %>% 
  group_by(treatment) %>%
  identify_outliers(max_peak)-> data_max_peak__nls_outliers

#Exclude outliers
data_max_peak_nls %>% 
  filter(!UI %in% data_max_peak__nls_outliers$UI) -> data_max_peak_nls_filt

# Build the linear model, excluding outliers
lm(max_peak ~ treatment*genotype, data = data_max_peak_nls_filt) -> model_filt 

 # Create a QQ plot of residuals and check for normality
ggqqplot(residuals(model_filt))
shapiro_test(residuals(model_filt))

##check Homogeneity of variance 
plot(model_filt, 1)
model_filt  %>% levene_test(max_peak ~ treatment*genotype)
##there is no homogeneity of variance, then we will apply a non-parametric test


### Kruskal Wallis on max peak
data_max_peak_nls_filt %>% 
   group_by(compartment, treatment) %>% 
  kruskal_test(max_peak~ genotype ) #%>% ####remove the hashtags if you want to create an excel file with the stats summary
 #write_xlsx(., path = "Krusk_MaxPeak_3.xlsx")

### Dunn's post-hoc test
data_max_peak_nls_filt %>% 
   group_by(compartment, treatment) %>% 
  dunn_test(max_peak~ genotype , p.adjust.method = "fdr")#%>% ####remove the hashtags if you want to create an excel file with the stats summary
## write_xlsx(., path = "Dunns_MaxPeak_3.xlsx")
  
```

###AUC_Figure 3
```{r} 
data_AUC_8  <- data_tidy %>% 
  filter  (time_after_treatment > 1) %>% 
  mutate(interval = case_when(time_after_treatment < 8 & time_after_treatment >1 ~ "Phase 1", 
                              #adding the factor" "interval" to indicate Phase 1 or Phase 2
                               time_after_treatment >= 8 ~ "Phase 2")) %>% 
  group_by(genotype, compartment, treatment, replicate, interval) %>% 
  summarise(AUC = sum(calcium)) 

data_AUC_8$interval <- as.factor(data_AUC_8$interval)

data_AUC_all  <- data_tidy %>% 
    filter  (time_after_treatment > 1) %>% 
  group_by(genotype, compartment, treatment, replicate) %>% 
  summarise(AUC = sum(calcium)) %>% 
  mutate (interval = "total") 

data_AUC_all$interval <- as.factor(data_AUC_all$interval)

data_AUC_merge <- rbind(data_AUC_all, data_AUC_8)

data_AUC_merge %>%   
filter(genotype!="cerk6")%>%
  filter(compartment=="nucleus")%>%
  split(.$treatment)%>%
  map(
~ggplot(., aes(x=genotype, y=AUC, fill=genotype))+
                scale_fill_manual(values=c("#cccccc", "#fe9929",  "#d95f0e")) +
                geom_dotplot(binaxis ="y", stackdir = "center", dotsize = 1.8) + 
                stat_summary(fun = median, fun.min = median, fun.max = median, geom = "crossbar", 
                           width = 0.7) +
                theme_classic()    +
  labs(y= expression(Ca^"2+"~integral~(mu*M))) +
    annotate ("text", label=paste0(unique(.$treatment)," ",unique(.$compartment)),  
              x=Inf, y=Inf,hjust=1,vjust=1, colour="black")+
  theme(axis.title.x=element_blank(), 
        strip.background = element_blank(),
        panel.border = element_rect(colour = "black", fill = NA))+
   facet_wrap(~interval)
) -> list_AUC_all

trace_nls[[1]]+
   list_max_all[[1]]+
   list_AUC_all[[1]]+
   trace_nls[[2]]+
   list_max_all[[2]]+
     list_AUC_all[[2]]+
   trace_nls[[3]]+
   list_max_all[[3]]+
     list_AUC_all[[3]]+
    plot_annotation(tag_levels = 'A') +  
  plot_layout(guides = 'collect', ncol = 3) -> plot_max_auc_nls_mutsymb
```

###check normality and homogeneity of variance on AUC_Figure 3
```{r}
data_AUC_merge %>% 
  filter(genotype!="cerk6") %>% 
  filter(compartment=="nucleus") %>%
mutate(UI = paste(compartment, treatment, genotype, replicate, AUC, interval, sep = '_')) -> data_AUC_nls

# Build the linear model
lm(AUC ~ genotype, data = data_AUC_nls) -> model_AUC 

 # Create a QQ plot of residuals and check for normality
ggqqplot(residuals(model_AUC))
shapiro_test(residuals(model_AUC))

##check Homogeneity of variance 
plot(model_AUC, 1)
model_AUC  %>% levene_test(AUC ~ genotype*treatment)
##there is no homogeneity of variance, then we will apply a non-parametric test

# Identify outliers
data_AUC_nls %>% 
  group_by(genotype, treatment) %>%
  identify_outliers(AUC)-> data_AUC_nls_outliers

#Exclude outliers
data_AUC_nls %>% 
  filter(!UI %in% data_AUC_nls_outliers$UI) -> data_AUC_nls_filt

# Build the linear model, excluding outliers
lm(AUC ~ genotype, data = data_AUC_nls_filt) -> model_AUC_nls_filt 

 # Create a QQ plot of residuals and check for normality
ggqqplot(residuals(model_AUC_nls_filt))
shapiro_test(residuals(model_AUC_nls_filt))

##check Homogeneity of variance 
plot(model_AUC_nls_filt, 1)
model_AUC_nls_filt  %>% levene_test(AUC ~ genotype*treatment)
##there is no homogeneity of variance, then we will apply a non-parametric test

### Kruskal Wallis on AUC
data_AUC_nls_filt %>% 
   group_by(compartment, interval, treatment) %>% 
  kruskal_test(AUC~ genotype) #%>% ####remove the hashtags if you want to create an excel file with the stats summary
 ##write_xlsx(., path = "Krusk_AUC_3.xlsx")

### Dunn's post-hoc test
data_AUC_nls_filt %>% 
   group_by(compartment, interval, treatment) %>% 
  dunn_test(AUC~ genotype , p.adjust.method = "fdr") #%>% ####remove the hashtags if you want to create an excel file with the stats summary
 ##write_xlsx(., path = "Dunns_AUC_3.xlsx")

```

###width_Figure 3
```{r}
### width > 0.4 totale
data_width_0.4_tot  <- data_tidy%>% 
  filter (time_after_treatment > 1) %>%  ###this parameter is to avoid touch
   mutate(over04 = case_when(calcium>0.4 ~ "true", 
                              #adding the threshold of calcium concentration 0.4
                              calcium<=0.4 ~ "false")) %>% 
       mutate(UI = paste(compartment, treatment, genotype, replicate, over04, sep = '_'))

data_width_0.4_tot_true <- data_width_0.4_tot %>%
  filter(over04=="true" )%>%
  group_by(genotype, compartment,  treatment, replicate) %>% 
   summarise(width_true = n()) %>% 
      mutate(UI = paste(compartment, treatment, genotype, replicate, sep = '_'))
  
data_width_0.4_tot_false <- data_width_0.4_tot %>%
  filter(over04=="false")%>%
  group_by(genotype, compartment,  treatment, replicate) %>% 
  summarise(width_false = n()) %>% 
        mutate(UI = paste(compartment, treatment, genotype, replicate, sep = '_'))

  
data_width_0.4_false2 <- anti_join(data_width_0.4_tot_false, data_width_0.4_tot_true)

data_width_0.4_total <- rbind(data_width_0.4_false2, data_width_0.4_tot_true)%>%
  select(., -c(width_false))
  
data_width_0.4_total[is.na(data_width_0.4_total)] <- 0

###plots with width
data_width_0.4_total %>%
  filter(genotype!="cerk6") %>% 
  filter(compartment=="nucleus") %>% 
  split(.$treatment) %>% 
  map(
~ggplot(., aes(x=genotype, y=width_true, fill=genotype))+
                scale_fill_manual(values=c("#cccccc", "#fe9929",  "#d95f0e")) +
                geom_dotplot(binaxis ="y", stackdir = "center", dotsize = 1.8) + 
                stat_summary(fun = median, fun.min = median, fun.max = median, geom = "crossbar", 
                           width = 0.7) +
                theme_classic()    +
labs(y=expression("width "*" [Ca"^"2+"*"] > 0.4 "*mu*M)) +
    annotate ("text", label=paste0(unique(.$treatment)," ",unique(.$compartment)),  
              x=Inf, y=Inf,hjust=1,vjust=1, colour="black")+
  theme(axis.title.x=element_blank(), 
        strip.background = element_blank(),
        panel.border = element_rect(colour = "black", fill = NA))
) -> list_width_0.4_total

trace_nls[[1]]+
list_width_0.4_total[[1]]+
trace_nls[[2]]+
list_width_0.4_total[[2]]+
  trace_nls[[3]]+
list_width_0.4_total[[3]]+
          plot_annotation(tag_levels = 'A') +  
  plot_layout(guides = 'collect', widths = c(2,0.8, 0.8),ncol = 3) -> plot_width_tot  
```

###check normality and homogeneity of variance on width 0.4_Figure 3
```{r}
data_width_0.4_total %>% 
  filter(genotype!="cerk6")%>%
  filter(compartment=="nucleus") %>% 
   mutate(UI = paste(genotype, compartment, treatment, replicate, sep = '_'))->data_width_0.4_nls

# Build the linear model
lm(width_true ~ genotype, data = data_width_0.4_nls) -> model_width_04

 # Create a QQ plot of residuals and check for normality
ggqqplot(residuals(model_width_04))
shapiro_test(residuals(model_width_04))

##check Homogeneity of variance 
plot(model_width_04, 1)
model_width_04  %>% levene_test(width_true ~ genotype)
##there is no homogeneity of variance, then we will apply a non-parametric test

# find the outliers
data_width_0.4_nls %>% 
  group_by(genotype) %>%
  identify_outliers(width_true)-> data_width_0.4_total_outliers

#exclude outliers
data_width_0.4_nls %>% 
  filter(!UI %in% data_width_0.4_total_outliers$UI) -> data_width_0.4_nls_filt

# Build the linear model, excluding outliers
lm(width_true ~ genotype, data = data_width_0.4_nls_filt) -> model_width_04_filt 

 # Create a QQ plot of residuals and check for normality
ggqqplot(residuals(model_width_04_filt))
shapiro_test(residuals(model_width_04_filt))

##check Homogeneity of variance 
plot(model_width_04_filt, 1)
model_width_04_filt  %>% levene_test(width_true ~ genotype)
##there is no homogeneity of variance, then we will apply a non-parametric test

### Kruskal Vallis on width
data_width_0.4_nls_filt %>% 
   group_by(compartment, treatment) %>% 
  kruskal_test(width_true~genotype ) #%>% ####remove the hashtags if you want to create an excel file with the stats summary
 ##write_xlsx(., path = "Krusk_width_3.xlsx")

### Dunn's post-hoc test
data_width_0.4_nls_filt %>% 
   group_by(compartment, treatment) %>% 
  dunn_test(width_true ~ genotype , p.adjust.method = "fdr") #%>% ####remove the hashtags if you want to create an excel file with the stats summary
 ##write_xlsx(., path = "Dunns_width_3.xlsx")
```
###plot for figure 3
```{r}
#####-------------Figure 3------------###########
trace_nls[[1]]+
  list_max_all[[1]]+
  list_AUC_all[[1]]+
  list_width_0.4_total[[1]]+
     trace_nls[[2]]+
  list_max_all[[2]]+
  list_AUC_all[[2]]+
  list_width_0.4_total[[2]]+
  trace_nls[[3]]+
  list_max_all[[3]]+
  list_AUC_all[[3]]+
  list_width_0.4_total[[3]]+
        plot_annotation(tag_levels = 'A') +  
  plot_layout(guides = 'collect', widths=c(1.3, 0.5, 1.5, 0.5), ncol = 4) #-> plot_fig3
```
###Figure 4

```{r}

###your path to the supplementary table "Dataset_S5.csv"
data <- read.csv("Dataset_S5_raw_data_Fig4.csv", header=T,sep = ";", na.string=c("na", "NA", "null")) 


```


###Figure 4 - Panel A
```{r}
#select only roi 6 root 6 from the dataset
roi6 <- data[ ,-(8:50)]
roi6 <- roi6[ ,-(2:6)]

# rmv NA prior the start of the recording
roi6 <- roi6[-(1:20),] 

roi6$timepoint <- as.numeric(roi6$timepoint)

#remove NA at the end of the trace
roi6 <- roi6%>%
  filter(roi6$timepoint<1830)

roi6 <- na.omit(roi6)

#tidy roi6
roi6 <- pivot_longer(roi6, cols=2, names_to= "sample", values_to = "ratio")

#create new time variable in minutes
roi6 <- roi6 %>%
  mutate(time_min = timepoint / 60)

roi6$time_min <- as.factor(roi6$time_min)


#graph root6 roi6 10-7M CO4 - panel A
ggplot (roi6, aes(time_min, ratio,   group=1)) +
  geom_line(lwd=1, color = "black") +
  scale_y_continuous(breaks=seq(0.75, 1.25, 0.2), expand = c(0,0), limits= c(0.75,1.25))+
  geom_vline(xintercept="8", linetype="dashed")+
  scale_x_discrete(roi6, breaks=c("10", "20", "30"))+
  theme_classic() +
  theme(axis.title.x=element_blank(),
        axis.text = element_text(size = 12),
        axis.title.y = element_blank()) -> plot_roi6_root6

```


###Figure 4 - Panel C
```{r}

#select only roi 2 root 8 from the dataset
data_2 <- data[ ,-(13:50)]
roi2 <- data_2[ ,-(2:11)]

# rmv NA prior the start of the recording
roi2 <- roi2[-(1:20),] 

roi2$timepoint <- as.numeric(roi2$timepoint)

#remove NA at the end of the trace
roi2 <- roi2%>%
  filter(roi2$timepoint<1830)

roi2 <- na.omit(roi2)

#tidy roi2
roi2 <- pivot_longer(roi2, cols=2, names_to= "sample", values_to = "ratio")

#create new time variable in minutes
roi2 <- roi2%>%
  mutate(time_min = timepoint / 60)

roi2$time_min <- as.factor(roi2$time_min)

# graph root8 roi2 10-7M CO4 - panel C
ggplot (roi2, aes(time_min, ratio,   group=1)) +
  geom_line(lwd=1, color = "black") +
  scale_y_continuous(breaks=seq(0.75, 1.25, 0.2), expand = c(0,0), limits= c(0.75,1.25))+
  geom_vline(xintercept="8", linetype="dashed")+
  scale_x_discrete(roi2, breaks=c("10", "20", "30"))+
  theme_classic() +
  theme(axis.title.x=element_blank(),
        axis.text = element_text(size = 12),
        axis.title.y = element_blank()) -> plot_roi2_root8

```

###Figure 4 - Panel B
```{r}
#select only roi 15 root 1 from the dataset
data_2 <- data[ ,-(38:50)]
roi15 <- data_2[ ,-(2:36)]

# rmv NA prior the start of the recording
roi15 <- roi15[-(1:20),] 

roi15$timepoint <- as.numeric(roi15$timepoint)

#remove NA at the end of the trace
roi15 <- roi15%>%
  filter(roi15$timepoint<1830)

roi15 <- na.omit(roi15)

#tidy roi15
roi15 <- pivot_longer(roi15, cols=2, names_to= "sample", values_to = "ratio")

#create new time variable in minutes
roi15 <- roi15%>%
  mutate(time_min = timepoint / 60)

roi15$time_min <- as.factor(roi15$time_min)

# graph root8 roi15 root 1 untreated - panel B
ggplot (roi15, aes(time_min, ratio,   group=1)) +
  geom_line(lwd=1, color = "black") +
  scale_y_continuous(breaks=seq(0.75, 1.25, 0.2), expand = c(0,0), limits= c(0.75,1.25))+
  geom_vline(xintercept="8", linetype="dashed")+
  scale_x_discrete(roi15, breaks=c("10", "20", "30"))+
  theme_classic() +
  theme(axis.title.x=element_blank(),
        axis.text = element_text(size = 12),
        axis.title.y = element_blank()) -> plot_roi15_root1_untreated
```

###Figure 4 - Panel D
```{r}

#select only roi 3 root 3 from the dataset
roi3 <- data[ ,-(41:50)]
roi3 <- roi3[ ,-(2:39)]

# rmv NA prior the start of the recording
roi3 <- roi3[-(1:20),] 

roi3$timepoint <- as.numeric(roi3$timepoint)

#remove NA at the end of the trace
roi3 <- roi3%>%
  filter(roi3$timepoint<1830)

roi3 <- na.omit(roi3)

#tidy roi3
roi3 <- pivot_longer(roi3, cols=2, names_to= "sample", values_to = "ratio")

#create new time variable in minutes
roi3 <- roi3%>%
  mutate(time_min = timepoint / 60)

roi3$time_min <- as.factor(roi3$time_min)

# graph root8 roi3 root 3 untreated - panel D
ggplot (roi3, aes(time_min, ratio,   group=1)) +
  geom_line(lwd=1, color = "black") +
  scale_y_continuous(breaks=seq(0.95, 1.45, 0.2), expand = c(0,0), limits= c(0.95,1.45))+
  geom_vline(xintercept="8", linetype="dashed")+
  scale_x_discrete(roi3, breaks=c("10", "20", "30"))+
  theme_classic() +
  theme(axis.title.x=element_blank(),
        axis.text = element_text(size = 12),
        axis.title.y = element_blank()) -> plot_roi3_root3_untreated

```

###Figure 4 - Panel E
```{r}

#select nuclei from root 6 only
root6 <- data[ ,-(11:50)]

# rmv NA prior the start of the recording
root6 <- root6[-(1:20),] 

root6$timepoint <- as.numeric(root6$timepoint)

#remove NA at the end of the trace
root6 <- root6%>%
  filter(root6$timepoint<1830)

root6 <- na.omit(root6)

#tidy root6
root6_tidy <- pivot_longer(root6, cols=2:10, names_to= "sample", values_to = "ratio")
root6_tidy <- root6_tidy %>% separate(sample, c('treatment', 'root', 'roi'))

root6_tidy <- root6_tidy %>%
  mutate(time_min = timepoint / 60)

root6_tidy$time_min <- as.factor(root6_tidy$time_min)
root6_tidy$roi <- as.factor(root6_tidy$roi)
root6_tidy$root <- as.factor(root6_tidy$root)
root6_tidy$timepoint <- as.factor(root6_tidy$timepoint)


###calculate mean and errors of root 6 by time
root6_stats  <- root6_tidy %>% 
  group_by(time_min) %>% 
  mutate(mean = mean(ratio), 
         stdev = sd(ratio),
         sterr = sd(ratio)/sqrt(length(ratio)),
         UI = paste(time_min, roi, sep = '_')) 

# graph mean root 6 10-7M - panel E
ggplot (root6_stats, aes(time_min, mean,   group=1)) +
  geom_line(lwd=1, color = "#7D7D7B") +
  geom_ribbon(aes(ymin=mean-sterr, ymax=mean+sterr),alpha=0.2,   colour = NA) +  
  scale_y_continuous(breaks=seq(0.75, 0.95, 0.05), expand = c(0,0), limits= c(0.75,0.95))+
  geom_vline(xintercept="8", linetype="dashed")+
  scale_x_discrete(root6_stats, breaks=c("10", "20", "30"))+
  theme_classic() +
  stat_smooth(color="#FF6A00", size = 1.5, method = "lm", formula = y ~ poly(x, 15))+
  theme(axis.title.x=element_blank(),
        axis.text = element_text(size = 12),
        axis.title.y = element_blank()) -> plot_mean_root6
```

###Figure 4 - Panel G
```{r}

#select nuclei from root 8 only
root8 <- data[ ,-(23:50)]
root8 <- root8[ ,-(2:10)]

# rmv NA prior the start of the recording
root8 <- root8[-(1:20),] 

root8$timepoint <- as.numeric(root8$timepoint)

#remove NA at the end of the trace
root8 <- root8%>%
  filter(root8$timepoint<1830)

root8 <- na.omit(root8)

#tidy root8
root8_tidy<- pivot_longer(root8, cols=2:12, names_to= "sample", values_to = "ratio")
root8_tidy <- root8_tidy %>% separate(sample, c('treatment', 'root', 'roi'))

root8_tidy <- root8_tidy %>%
  mutate(time_min = timepoint / 60)

root8_tidy$time_min <- as.factor(root8_tidy$time_min)
root8_tidy$roi <- as.factor(root8_tidy$roi)
root8_tidy$root <- as.factor(root8_tidy$root)
root8_tidy$timepoint <- as.factor(root8_tidy$timepoint)


###calculate mean and errors of root by time
root8_stats  <- root8_tidy %>% 
  group_by(time_min) %>% 
  mutate(mean = mean(ratio), 
         stdev = sd(ratio),
         sterr = sd(ratio)/sqrt(length(ratio)),
         UI = paste(time_min, roi, sep = '_')) 


# graph mean root 8 10-7M - panel G
ggplot (root8_stats, aes(time_min, mean,   group=1)) +
  geom_line(lwd=1, color = "#7D7D7B") +
  geom_ribbon(aes(ymin=mean-sterr, ymax=mean+sterr),alpha=0.2,   colour = NA) +  
  scale_y_continuous(breaks=seq(0.8, 1, 0.05), expand = c(0,0), limits= c(0.8,1))+
  geom_vline(xintercept="8", linetype="dashed")+
  scale_x_discrete(root8_stats, breaks=c("10", "20", "30"))+
  theme_classic()+
  stat_smooth(color="#FF6A00", size = 1.5, method = "lm", formula = y ~ poly(x, 15))+
theme(axis.title.x=element_blank(),
      axis.text = element_text(size = 12),
      axis.title.y = element_blank()) -> plot_mean_root8
```

###Figure 4 - Panel F
```{r}

#select nuclei from root 1 only
root1 <- data[ ,-(38:50)]
root1 <- root1[ ,-(2:22)]

# rmv NA prior the start of the recording
root1 <- root1[-(1:20),] 

root1$timepoint <- as.numeric(root1$timepoint)

#remove NA at the end of the trace
root1 <- root1%>%
  filter(root1$timepoint<1830)

root1 <- na.omit(root1)

#tidy root1
root1_tidy<- pivot_longer(root1, cols=2:16, names_to= "sample", values_to = "ratio")
root1_tidy <- root1_tidy %>% separate(sample, c('treatment', 'root', 'roi'))

root1_tidy <- root1_tidy %>%
  mutate(time_min = timepoint / 60)

root1_tidy$time_min <- as.factor(root1_tidy$time_min)
root1_tidy$roi <- as.factor(root1_tidy$roi)
root1_tidy$root <- as.factor(root1_tidy$root)
root1_tidy$timepoint <- as.factor(root1_tidy$timepoint)


###calculate mean and errors of root by time
root1_stats  <- root1_tidy %>% 
  group_by(time_min) %>% 
  mutate(mean = mean(ratio), 
         stdev = sd(ratio),
         sterr = sd(ratio)/sqrt(length(ratio)),
         UI = paste(time_min, roi, sep = '_')) 

# graph mean root 1 untreated - panel F
ggplot (root1_stats, aes(time_min, mean,   group=1)) +
  geom_line(lwd=1, color = "#7D7D7B") +
  geom_ribbon(aes(ymin=mean-sterr, ymax=mean+sterr),alpha=0.2,   colour = NA) +  
  scale_y_continuous(breaks=seq(0.92, 1.12, 0.05), expand = c(0,0), limits= c(0.92,1.12))+
  geom_vline(xintercept="8", linetype="dashed")+
  scale_x_discrete(root1_stats, breaks=c("10", "20", "30"))+
  stat_smooth(color="#FF6A00", size = 1.5, method = "lm", formula = y ~ poly(x, 15))+
  theme_classic() +
  theme(axis.title.x=element_blank(),
        axis.text = element_text(size = 12),
        axis.title.y = element_blank()) -> plot_mean_root1_untreated


```

###Figure 4 - Panel H
```{r}
#select nuclei from root 3 only
root3 <- data[ ,-(2:37)]

# rmv NA prior the start of the recording
root3 <- root3[-(1:20),] 

root3$timepoint <- as.numeric(root3$timepoint)

#remove NA at the end of the trace
root3 <- root3%>%
  filter(root3$timepoint<1830)

root3 <- na.omit(root3)

#tidy root3
root3_tidy<- pivot_longer(root3, cols=2:14, names_to= "sample", values_to = "ratio")
root3_tidy <- root3_tidy %>% separate(sample, c('treatment', 'root', 'roi'))

root3_tidy <- root3_tidy %>%
  mutate(time_min = timepoint / 60)

root3_tidy$time_min <- as.factor(root3_tidy$time_min)
root3_tidy$roi <- as.factor(root3_tidy$roi)
root3_tidy$root <- as.factor(root3_tidy$root)
root3_tidy$timepoint <- as.factor(root3_tidy$timepoint)


###calculate mean and errors of root by time
root3_stats<-root3_tidy%>%
  group_by(time_min) %>% 
  mutate(mean = mean(ratio), 
         stdev = sd(ratio),
         sterr = sd(ratio)/sqrt(length(ratio)),
         UI = paste(time_min, roi, sep = '_')) 

# graph mean root 3 untreated - panel H
ggplot (root3_stats, aes(time_min, mean,   group=1)) +
  geom_line(lwd=1, color = "#7D7D7B") +
  geom_ribbon(aes(ymin=mean-sterr, ymax=mean+sterr),alpha=0.2,   colour = NA) +  
  scale_y_continuous(breaks=seq(1.1, 1.3, 0.05), expand = c(0,0), limits= c(1.1,1.3))+
  geom_vline(xintercept="8", linetype="dashed")+
  scale_x_discrete(root3_stats, breaks=c("10", "20", "30"))+
  stat_smooth(color="#FF6A00", size = 1.5, method = "lm", formula = y ~ poly(x, 15))+
  theme_classic() +
  theme(axis.title.x=element_blank(),
        axis.text = element_text(size = 12),
        axis.title.y = element_blank()) -> plot_mean_root3_untreated

```

###Assemble panels into Fig. 4
```{r}
#######--------Figure 4----------########
plot_roi6_root6+
  plot_roi15_root1_untreated+
  plot_roi2_root8+
  plot_roi3_root3_untreated+
  plot_mean_root6+
  plot_mean_root1_untreated+
  plot_mean_root8+
  plot_mean_root3_untreated+
  plot_annotation(tag_levels = 'A') +  
  plot_layout(guides = 'collect', heights = c(1, 1, 1.7, 1.7), ncol = 2) #-> plot_fig4
```

###Figure 5
###Loading and tidying aequorin files_Figure 5
```{r}
###your path to the supplementary table "Dataset_S4_raw_data_Fig3-5-S5.csv"
data <- read.csv("Dataset_S4_raw_data_Fig3-5-S5.csv", 
                 header=T,sep = ";", na.string=c("na", "NA", "null")) 
str(data)
data_tidy <- pivot_longer(data, cols=2:ncol(data), names_to= "condition", values_to = "calcium")

data_tidy <- data_tidy %>% separate(condition, c('compartment', 'treatment','genotype', 'replicate'))

data_tidy$compartment <- as.factor(data_tidy$compartment)
data_tidy$treatment <- as.factor(data_tidy$treatment)
data_tidy$genotype <- as.factor(data_tidy$genotype)
data_tidy$replicate <- as.factor(data_tidy$replicate)
data_tidy$genotype <- factor (data_tidy$genotype, levels = c("wt", "symrk", "castor", "cerk6"))

data_tidy$treatment <- recode_factor(data_tidy$treatment, co4 = "CO4", co8 = "CO8", mycLCO = "mycLCO")
data_tidy$compartment <- recode_factor(data_tidy$compartment, cya="cytosol", nls="nucleus")

data_tidy <- data_tidy %>% 
  droplevels()

data_tidy <- na.omit(data_tidy) 
head(data_tidy)

data_tidy <- data_tidy %>% 
  mutate (time_after_treatment = time - 1.667)
```

###new dataset with stats: mean, standard deviation, standard error_Figure 5
```{r}
data_stats  <- data_tidy %>% 
  group_by(time_after_treatment, genotype, compartment, treatment) %>% 
  summarise(mean = mean(calcium), 
            stdev = sd(calcium),
            sterr = sd(calcium)/sqrt(length(calcium))
             ) 
```

###panel A (CO4 cytosolic)_Figure 5
```{r}

###new dataset for CO4-induced cytosolic response in cerk6 mutant
data_stats_cerk6_cya_CO4  <- data_stats %>% 
      filter(genotype == "cerk6")%>%
   filter(compartment == "cytosol")%>%
  filter(treatment == "CO4")
  
###calculation of rolling average for CO4-induced cytosolic response in wt and symrk mutants
data_rollavg_solomean_wt_cya_CO4  <-data_stats%>%
    filter(genotype == "wt")%>%
  filter(compartment == "cytosol")%>%
  filter(treatment == "CO4")%>%
  group_by(genotype, compartment, treatment) %>% 
  mutate(rollavg_9 = zoo::rollmean(mean, 9, fill=NA)
         )

data_rollavg_solomean_symrk_cya_CO4  <-data_stats %>% 
    filter(genotype == "symrk")%>%
  filter(compartment == "cytosol")%>%
  filter(treatment == "CO4")%>%
  group_by(genotype, compartment, treatment) %>% 
  mutate(rollavg_9 = zoo::rollmean(mean, 9, fill=NA)
         )
### panel A
ggplot (data_stats_cerk6_cya_CO4, aes(time_after_treatment, mean, color="#998ec3")) +
  geom_line(lwd=1) +
    scale_colour_manual(values=c("#998ec3")) + 
  scale_fill_manual(values=c("#998ec3")) +
   geom_ribbon(aes(ymin=mean-sterr, ymax=mean+sterr),alpha=0.2,color=NA, fill="#998ec3") +  
  #scale_y_continuous(breaks=seq(0, 1.5, 0.2), expand = c(0,0), limits= c(0,1.5))+
 #ylim (0, 1.25) +
  scale_x_continuous(expand = c(0,0))+
 geom_line(data = data_rollavg_solomean_wt_cya_CO4, aes(time_after_treatment, rollavg_9), color="#cccccc", lwd=1.2, linetype="dashed" )+
 geom_line(data = data_rollavg_solomean_symrk_cya_CO4, aes(time_after_treatment, rollavg_9), color= "#f1a340", lwd=1.2, linetype="dashed" )+ 
   theme_classic() +
  annotate("segment", x = 8, xend = 8, y = 0, yend = Inf, colour="#737373",linetype = "dashed") +
  annotate("segment", x = 0, y = 0.9, xend = 0, yend = 0.8,lineend = "round", linejoin = "round",
         arrow = arrow(type = "closed", length = unit(0.02, "npc"))) +
    annotate ("text", x=5, y=1.6, label="Phase 1", colour="#737373")+
annotate ("text", x=15, y=1.6, label="Phase 2", colour="#737373")+
   annotate ("text", x=0, y=1, label="CO4", colour="black")+
  labs(y="Cytosolic Ca^(2+) conc (\u03bcM)") + # \u03bc is the Unicode for Greek mu
  #labs(y=paste0(unique(.$compartment)," Ca2+ conc")) +
    theme(axis.title.x=element_blank()) -> trace_cya_CO4
```


###panel C (CO8 cytosolic)_Figure 5
```{r}
###new dataset for CO8-induced cytosolic response in cerk6 mutant
data_stats_cerk6_cya_CO8  <- data_stats %>% 
      filter(genotype == "cerk6")%>%
   filter(compartment == "cytosol")%>%
  filter(treatment == "CO8")
  
###calculation of rolling average for CO8-induced cytosolic response in wt and symrk mutants
data_rollavg_solomean_wt_cya_CO8  <-data_stats%>%
    filter(genotype == "wt")%>%
  filter(compartment == "cytosol")%>%
  filter(treatment == "CO8")%>%
  group_by(genotype, compartment, treatment) %>% 
  mutate(rollavg_9 = zoo::rollmean(mean, 9, fill=NA)
         )

data_rollavg_solomean_symrk_cya_CO8  <-data_stats %>% 
    filter(genotype == "symrk")%>%
  filter(compartment == "cytosol")%>%
  filter(treatment == "CO8")%>%
  group_by(genotype, compartment, treatment) %>% 
  mutate(rollavg_9 = zoo::rollmean(mean, 9, fill=NA))

### panel C
ggplot (data_stats_cerk6_cya_CO8, aes(time_after_treatment, mean, color="#998ec3")) +
  geom_line(lwd=1) +
    scale_colour_manual(values=c("#998ec3")) + 
  scale_fill_manual(values=c("#998ec3")) +
   geom_ribbon(aes(ymin=mean-sterr, ymax=mean+sterr),alpha=0.2,color=NA, fill="#998ec3") +  
  #scale_y_continuous(breaks=seq(0, 1.5, 0.2), expand = c(0,0), limits= c(0,1.5))+
 #ylim (0, 1.25) +
  scale_x_continuous(expand = c(0,0))+
 geom_line(data = data_rollavg_solomean_wt_cya_CO8, aes(time_after_treatment, rollavg_9), color="#cccccc", lwd=1.2, linetype="dashed" )+
 geom_line(data = data_rollavg_solomean_symrk_cya_CO8, aes(time_after_treatment, rollavg_9), color= "#f1a340", lwd=1.2, linetype="dashed" )+ 
   theme_classic() +
  annotate("segment", x = 8, xend = 8, y = 0, yend = Inf, colour="#737373",linetype = "dashed") +
  annotate("segment", x = 0, y = 0.9, xend = 0, yend = 0.8,lineend = "round", linejoin = "round",
         arrow = arrow(type = "closed", length = unit(0.02, "npc"))) +
    annotate ("text", x=5, y=1.6, label="Phase 1", colour="#737373")+
annotate ("text", x=15, y=1.6, label="Phase 2", colour="#737373")+
   annotate ("text", x=0, y=1, label="CO8", colour="black")+
  labs(y="Cytosolic Ca^(2+) conc (\u03bcM)") + # \u03bc is the Unicode for Greek mu
  #labs(y=paste0(unique(.$compartment)," Ca2+ conc")) +
    theme(axis.title.x=element_blank()) -> trace_cya_CO8
```

###panel B (CO4 nuclear)_Figure 5
```{r}

###new dataset for CO4-induced nuclear response in cerk6 mutant
data_stats_cerk6_nls_CO4  <- data_stats %>% 
      filter(genotype == "cerk6")%>%
   filter(compartment == "nucleus")%>%
  filter(treatment == "CO4")
  
###calculation of rolling average for CO4-induced nuclear response in wt and symrk mutants
data_rollavg_solomean_wt_nls_CO4  <-data_stats%>%
    filter(genotype == "wt")%>%
  filter(compartment == "nucleus")%>%
  filter(treatment == "CO4")%>%
  group_by(genotype, compartment, treatment) %>% 
  mutate(rollavg_9 = zoo::rollmean(mean, 9, fill=NA)
         )

data_rollavg_solomean_symrk_nls_CO4  <-data_stats %>% 
    filter(genotype == "symrk")%>%
  filter(compartment == "nucleus")%>%
  filter(treatment == "CO4")%>%
  group_by(genotype, compartment, treatment) %>% 
  mutate(rollavg_9 = zoo::rollmean(mean, 9, fill=NA))

### panel B
ggplot (data_stats_cerk6_nls_CO4, aes(time_after_treatment, mean, color="#998ec3")) +
  geom_line(lwd=1) +
    scale_colour_manual(values=c("#998ec3")) + 
  scale_fill_manual(values=c("#998ec3")) +
   geom_ribbon(aes(ymin=mean-sterr, ymax=mean+sterr),alpha=0.2,color=NA, fill="#998ec3") +  
  #scale_y_continuous(breaks=seq(0, 1.5, 0.2), expand = c(0,0), limits= c(0,1.5))+
 #ylim (0, 1.25) +
  scale_x_continuous(expand = c(0,0))+
 geom_line(data = data_rollavg_solomean_wt_nls_CO4, aes(time_after_treatment, rollavg_9), color="#cccccc", lwd=1.2, linetype="dashed" )+
 geom_line(data = data_rollavg_solomean_symrk_nls_CO4, aes(time_after_treatment, rollavg_9), color= "#f1a340", lwd=1.2, linetype="dashed" )+ 
   theme_classic() +
  annotate("segment", x = 8, xend = 8, y = 0, yend = Inf, colour="#737373",linetype = "dashed") +
  annotate("segment", x = 0, y = 0.9, xend = 0, yend = 0.8,lineend = "round", linejoin = "round",
         arrow = arrow(type = "closed", length = unit(0.02, "npc"))) +
    annotate ("text", x=5, y=1.6, label="Phase 1", colour="#737373")+
annotate ("text", x=15, y=1.6, label="Phase 2", colour="#737373")+
   annotate ("text", x=0, y=1, label="CO4", colour="black")+
  labs(y="nuclear Ca^(2+) conc (\u03bcM)") + # \u03bc is the Unicode for Greek mu
     theme(axis.title.x=element_blank()) -> trace_nls_CO4

```


###panel D (CO8 nuclear)_Figure 5
```{r}
###new dataset for CO8-induced nuclear response in cerk6 mutant
data_stats_cerk6_nls_CO8  <- data_stats %>% 
      filter(genotype == "cerk6")%>%
   filter(compartment == "nucleus")%>%
  filter(treatment == "CO8")
  
###calculation of rolling average for CO8-induced nuclear response in wt and symrk mutants
data_rollavg_solomean_wt_nls_CO8  <-data_stats%>%
    filter(genotype == "wt")%>%
  filter(compartment == "nucleus")%>%
  filter(treatment == "CO8")%>%
  group_by(genotype, compartment, treatment) %>% 
  mutate(rollavg_9 = zoo::rollmean(mean, 9, fill=NA)
         )

data_rollavg_solomean_symrk_nls_CO8  <-data_stats %>% 
    filter(genotype == "symrk")%>%
  filter(compartment == "nucleus")%>%
  filter(treatment == "CO8")%>%
  group_by(genotype, compartment, treatment) %>% 
  mutate(rollavg_9 = zoo::rollmean(mean, 9, fill=NA))

### panel D
ggplot (data_stats_cerk6_nls_CO8, aes(time_after_treatment, mean, color="#998ec3")) +
  geom_line(lwd=1) +
    scale_colour_manual(values=c("#998ec3")) + 
  scale_fill_manual(values=c("#998ec3")) +
   geom_ribbon(aes(ymin=mean-sterr, ymax=mean+sterr),alpha=0.2,color=NA, fill="#998ec3") +  
  #scale_y_continuous(breaks=seq(0, 1.5, 0.2), expand = c(0,0), limits= c(0,1.5))+
 #ylim (0, 1.25) +
  scale_x_continuous(expand = c(0,0))+
 geom_line(data = data_rollavg_solomean_wt_nls_CO8, aes(time_after_treatment, rollavg_9), color="#cccccc", lwd=1.2, linetype="dashed" )+
 geom_line(data = data_rollavg_solomean_symrk_nls_CO8, aes(time_after_treatment, rollavg_9), color= "#f1a340", lwd=1.2, linetype="dashed" )+ 
   theme_classic() +
  annotate("segment", x = 8, xend = 8, y = 0, yend = Inf, colour="#737373",linetype = "dashed") +
  annotate("segment", x = 0, y = 0.9, xend = 0, yend = 0.8,lineend = "round", linejoin = "round",
         arrow = arrow(type = "closed", length = unit(0.02, "npc"))) +
    annotate ("text", x=5, y=1.6, label="Phase 1", colour="#737373")+
annotate ("text", x=15, y=1.6, label="Phase 2", colour="#737373")+
   annotate ("text", x=0, y=1, label="CO8", colour="black")+
  labs(y="nuclear Ca^(2+) conc (\u03bcM)") + # \u03bc is the Unicode for Greek mu
     theme(axis.title.x=element_blank()) -> trace_nls_CO8

```

####final plot fig.5
```{r}
#######--------Figure 5---------########
  trace_cya_CO4+ 
    trace_nls_CO4+
    trace_cya_CO8 + 
    trace_nls_CO8 +
   plot_annotation(tag_levels = 'A') +  
  plot_layout(guides = 'collect', ncol = 2)# -> plot_Fig5
```


###Figure 6
##Loading and tidying aequorin files_Figure 6
```{r}
###your path to the supplementary table "Dataset_S6_raw_data_Fig6A-D.csv"

data <- read.csv("Dataset_S6_raw_data_Fig6A-D_S3.csv", 
                 header=T,sep = ";", na.string=c("na", "NA", "null")) 
str(data)
data_tidy <- pivot_longer(data, cols=2:ncol(data), names_to= "condition", values_to = "calcium")

data_tidy <- data_tidy %>% separate(condition, c('compartment', 'treatment',
                                                 'dilution', 'replicate'))

data_tidy$compartment <- as.factor(data_tidy$compartment)
data_tidy$treatment <- as.factor(data_tidy$treatment)
data_tidy$dilution <- as.factor(data_tidy$dilution)
data_tidy$replicate <- as.factor(data_tidy$replicate)

data_tidy$treatment <- recode_factor(data_tidy$treatment, co4 = "CO4", 
                                     co8 = "CO8")
data_tidy$compartment <- recode_factor(data_tidy$compartment, cya = "cytosol",nls = "nucleus")

data_tidy$dilution <- factor(data_tidy$dilution, levels=c('6','5x7', '7','8', '9'))

data_tidy <- na.omit(data_tidy) 
head(data_tidy)

data_tidy <- data_tidy %>% 
  mutate (time_after_treatment=data_tidy$`time.point`-1.667)
```

###new dataset with stats: mean, standard deviation, standard error_Figure 6
```{r}
data_stats  <- data_tidy %>% 
  group_by(time_after_treatment, compartment, dilution, treatment) %>% 
  summarise(mean = mean(calcium), 
            stdev = sd(calcium),
            sterr = sd(calcium)/sqrt(length(calcium))
             ) 
```

####plot_aequorin_trace_CO4_Figure 6
```{r}
data_stats %>% 
  filter(treatment=="CO4") %>% 
  split(., list(.$compartment)) %>% 
  map(
~ggplot (., aes(time_after_treatment, mean,   color=dilution, fill=dilution)) +
  geom_line(lwd=1) +
  scale_colour_manual(values=c("#252525","#636363", "#969696", "#cccccc")) + 
  scale_fill_manual(values=c("#252525","#636363", "#969696", "#cccccc")) +
   geom_ribbon(aes(ymin=mean-sterr, ymax=mean+sterr),alpha=0.25,   colour = NA) +   
  scale_y_continuous(breaks=seq(0, 1.8, 0.2), expand = c(0,0), limits= c(0,1.8))+
  scale_x_continuous("time (min)", expand = c(0,0), limits = c(-1.67,29.5))+
  theme_classic(base_size = 12)  +
  annotate("segment", x = 8, xend = 8, y = 0, yend = Inf, colour="#737373",linetype = "dashed") +
  annotate("segment", x = 0, y = 1.65, xend = 0, yend = 1.42,lineend = "round", linejoin = "round",
         arrow = arrow(type = "closed", length = unit(0.02, "npc"))) +
#  annotate ("text", x=5, y=1.05, label="Phase 1", colour="#737373", size= 7)+
 # annotate ("text", x=15, y=1.05, label="Phase 2", colour="#737373", size= 7)+
    annotate ("text", x=-0.6, y=1.73, label="CO4", colour="black", size= 4, hjust = 0)+
  #labs(y="Nuclear Ca^(2+) conc (\u03bcM)") + # \u03bc is the Unicode for Greek mu
 #labs(y=paste0(unique(.$compartment)," Ca2+ conc") +
 labs(y= expression(Ca^"2+"~conc~(mu*M)), x=expression (Time~"(min)")) +
  theme(strip.background = element_blank()) 
 )-> trace_CO4
```


####plot_aequorin_trace_CO8_Figure S4
```{r}
data_stats %>% 
  filter(treatment=="CO8") %>% 
  ggplot (., aes(time_after_treatment, mean,   color=dilution, fill=dilution)) +
  geom_line(lwd=1) +
  scale_colour_manual(values=c("#252525", "#636363", "#969696")) + 
  scale_fill_manual(values=c("#252525", "#636363", "#969696")) +
   geom_ribbon(aes(ymin=mean-sterr, ymax=mean+sterr),alpha=0.25,   colour = NA) +   
  scale_y_continuous(breaks=seq(0, 1.8, 0.2), expand = c(0,0), limits= c(0,1.8))+
  scale_x_continuous("time (min)", expand = c(0,0), limits = c(-1.67,29.5))+
  theme_classic(base_size = 12)  +
  annotate("segment", x = 8, xend = 8, y = 0, yend = Inf, colour="#737373",linetype = "dashed") +
  annotate("segment", x = 0, y = 1.65, xend = 0, yend = 1.42,lineend = "round", linejoin = "round",
         arrow = arrow(type = "closed", length = unit(0.02, "npc"))) +
#  annotate ("text", x=5, y=1.05, label="Phase 1", colour="#737373", size= 7)+
 # annotate ("text", x=15, y=1.05, label="Phase 2", colour="#737373", size= 7)+
    annotate ("text", x=-0.6, y=1.73, label="CO8", colour="black", size= 4, hjust = 0)+
  #labs(y="Nuclear Ca^(2+) conc (\u03bcM)") + # \u03bc is the Unicode for Greek mu
 #labs(y=paste0(unique(.$compartment)," Ca2+ conc") +
 labs(y= expression(Ca^"2+"~conc~(mu*M)), x=expression (Time~"(min)")) +
  theme(strip.background = element_blank()) +
  facet_wrap(~compartment)-> trace_CO8
```


##Area Under the Curve (AUC) divided by phases_Figure 6
```{r}
data_AUC_8  <- data_tidy %>% 
    filter  (time_after_treatment > 1.5 & time_after_treatment < 29.5) %>% 
  mutate(interval = case_when(time_after_treatment < 8 & time_after_treatment >1 ~ "Phase 1", 
                              #adding the factor" "interval" to indicate Phase 1 or Phase 2
                               time_after_treatment >= 8 ~ "Phase 2")) %>% 
  group_by(dilution, compartment, treatment, replicate, interval) %>% 
  summarise(AUC = sum(calcium)) 

data_AUC_8$interval <- as.factor(data_AUC_8$interval)
```


##total AUC _Figure 6
```{r}
data_AUC_all  <- data_tidy %>% 
  group_by(dilution, compartment, treatment, replicate) %>% 
  summarise(AUC = sum(calcium)) %>% 
  mutate (interval = "total") 
data_AUC_all$interval <- as.factor(data_AUC_all$interval)
```
##merge AUC data_Figure 6
```{r}
data_AUC_merge <- rbind(data_AUC_all, data_AUC_8)

### Kruskal Wallis on max peak
data_AUC_merge %>% 
  filter (interval != "total") %>% 
   group_by(compartment, treatment, interval) %>% 
  kruskal_test(AUC~ dilution ) #%>% ####remove the hashtags if you want to create an excel file with the stats summary
 #write_xlsx(., path = "Krusk_Max_AUC_5.xlsx")

### Dunn's post-hoc test
data_AUC_merge %>% 
   filter (interval != "total") %>% 
   group_by(compartment, treatment, interval) %>% 
  dunn_test(AUC~ dilution , p.adjust.method = "fdr") #%>% ####remove the hashtags if you want to create an excel file with the stats summary
 #write_xlsx(., path = "Dunns_Max_AUC_5.xlsx")
```

##dotplot AUC, CO4_Figure 6
```{r}
data_AUC_8 %>% 
  filter (treatment=="CO4") %>% 
  split(., list(.$compartment)) %>% 
  map(
~ggplot (., aes(x=dilution, y=AUC,   fill=dilution)) +
              scale_fill_manual(values=c("#252525","#636363", "#969696", "#cccccc")) + 
              geom_dotplot(binaxis ="y", stackdir = "center", dotsize = 1.2) + 
              stat_summary(fun = median, fun.min = median, fun.max = median, 
                           geom = "crossbar", width = 0.7) +
              theme_classic(base_size = 12)    +
  labs(y= expression(Integral~Ca^"2+"~(mu*M)), x=expression(dilution)) +
    annotate ("text", label=paste0("CO4",unique(.$compartment)),
              x=0.5, y=9, colour="black", hjust = 0)+
  theme(strip.background = element_blank())+
        facet_wrap(~interval)
)-> list_AUC_8_CO4
```

##dotplot AUC, CO8_Figure S3
```{r}
data_AUC_8 %>% 
  filter (treatment=="CO8") %>% 
  split(., list(.$compartment)) %>% 
  map(
~ggplot (., aes(x=dilution, y=AUC,   fill=dilution)) +
              scale_fill_manual(values=c("#252525", "#636363", "#969696")) + 
              geom_dotplot(binaxis ="y", stackdir = "center", dotsize = 1.2) + 
              stat_summary(fun = median, fun.min = median, fun.max = median, 
                           geom = "crossbar", width = 0.7) +
              theme_classic(base_size = 12)    +
  labs(y= expression(Integral~Ca^"2+"~(mu*M)), x=expression(dilution)) +
    annotate ("text", label=paste0("CO8",unique(.$compartment)),
              x=0.5, y=9, colour="black", hjust = 0)+
  theme(strip.background = element_blank())+
        facet_wrap(~interval)
)-> list_AUC_8_CO8
```

##arrange plots with only phases_Figure 6
```{r}
######-------Figure6A-D------#####
  trace_CO4[[1]] + 
  list_AUC_8_CO4[[1]] +
   trace_CO4[[2]] +
  list_AUC_8_CO4[[2]] +
   plot_annotation(tag_levels = 'A') +  
  plot_layout(guides = 'collect', widths = c(1.5,1,1.5,1),nrow = 1) #->plot_Fig6AD
```

###Figure 6 panel E

###Loading and tidying qPCR data
```{r}
data <- read.csv("Dataset_S8_raw_data_Fig6E.csv", 
                 header=T,sep = ";", na.string=c("na", "NA", "null")) 
str(data)

data$gene <- as.factor(data$gene)
data$genotype <- as.factor(data$genotype)
data$treatment <- factor(data$treatment, levels=c("10-7 CO4", "10-9 CO4","ctr"))
data$replicate <- as.factor(data$replicate)


```

###plot ratio - 1h_Figure 6
```{r} 
data%>%
  split(.$gene)%>%
map(
~ggplot(., aes(x=treatment, y=Ratio, fill=treatment))+
                scale_fill_manual(values=c("#636363","#cccccc", "#f7f7f7")) +
                #scale_color_manual(values=c("#636363","#bdbdbd", "#f0f0f0")) +
                geom_dotplot(binaxis ="y", stackdir = "center", dotsize = 1.8) + 
                stat_summary(fun = median, fun.min = median, fun.max = median, geom = "crossbar", 
                           width = 0.6) +
                theme_classic()    +
  labs(title =paste0(unique(.$gene)))+
  labs(y= expression("Arbitrary expression unit")) +
  theme(axis.title.x=element_blank(), 
        strip.background = element_blank(),
        panel.border = element_rect(colour = "black", fill = NA))
) -> plot

######-------Figure6E------#####
plot[[1]]+
  plot[[3]]+
  plot[[4]]+
  plot[[2]]+
    plot_annotation(tag_levels = 'A') +  
  plot_layout(guides = 'collect', nrow = 1) ##-> plot_6E

```

###check normality and homogeneity of variance on ratio - chitinase_Figure 6
```{r}
data%>%
  filter(genotype=="WT")%>%
   filter(gene=="chitinase")  -> data_chitinase

# Build the linear model
lm(Ratio ~ treatment, data = data_chitinase) -> model

# Create a QQ plot of residuals and check for normality
ggqqplot(residuals(model))
shapiro_test(residuals(model))

##check Homogeneity of variance 
plot(model, 1)
model  %>% levene_test(Ratio ~ treatment)
##there is  homogeneity of variance, then we will apply Anova

```

###check normality and homogeneity of variance on ratio - Rbohb
```{r}

data%>%
   filter(gene=="RbohB-like")  -> data_Rbohb

# Build the linear model
lm(Ratio ~ treatment, data = data_Rbohb) -> model

# Create a QQ plot of residuals and check for normality
ggqqplot(residuals(model))
shapiro_test(residuals(model))

##check Homogeneity of variance 
plot(model, 1)
model  %>% levene_test(Ratio ~ treatment)
##there is homogeneity of variance, then we will apply Anova
```

###check normality and homogeneity of variance on ratio - WRKY70-like
```{r}

data%>%
   filter(gene=="WRKY70-like")  -> data_WRKY70

# Build the linear model
lm(Ratio ~ treatment, data = data_WRKY70) -> model

# Create a QQ plot of residuals and check for normality
ggqqplot(residuals(model))
shapiro_test(residuals(model))

##check Homogeneity of variance 
plot(model, 1)
model  %>% levene_test(Ratio ~ treatment)
##there is homogeneity of variance, then we will apply Anova
```

###check normality and homogeneity of variance on ratio - PRp27-like
```{r}
data%>%
  filter(genotype=="WT")%>%
   filter(gene=="PRp27-like") %>%
  mutate(UI = paste(gene, genotype, treatment, replicate, sep = '_'))-> data_PRp27

# Build the linear model
lm(Ratio ~ treatment, data = data_PRp27) -> model

# Create a QQ plot of residuals and check for normality
ggqqplot(residuals(model))
shapiro_test(residuals(model))

##check Homogeneity of variance 
plot(model, 1)
model  %>% levene_test(Ratio ~ treatment)
##there is homogeneity of variance, then we will apply Anova
```

###Anova and Tukey on qPCR data_Figure 6E
```{r}
### ANOVA
data %>%  split(., list(.$gene)) %>%
  map (
    ~anova_test (., Ratio ~ treatment)
  ) #%>% ####remove the hashtags if you want to create an excel file with the stats summary
# write_xlsx(., path = "Anova_qPCR.xlsx")


### Tukey
data %>%  split(., list(.$gene)) %>%
  map (
    ~tukey_hsd (., Ratio ~ treatment)
  ) #%>% ####remove the hashtags if you want to create an excel file with the stats summary
 #write_xlsx(., path = "Tukey_qPCR.xlsx")
```


###Supplementary Figure S4

```{r}
###your path to the supplementary Dataset S7
data <-read.csv("Dataset_S7_raw_data_FigS4.csv", 
                   header=T,sep = ";", na.string=c("na", "NA", "null")) 
```

###Supplementary Fig S4 - Panel A
```{r}
#select only roi 7 root 2 from the dataset
roi7 <- data[,-(7:20)]
roi7 <- roi7[,-(2:5)]

# rmv NA prior the start of the recording
roi7 <- roi7[-(1:17),] 

#remove NA at the end of the trace
roi7$timepoint <- as.numeric(roi7$timepoint)
roi7 <- roi7%>%
  filter(roi7$timepoint<1835)

roi7 <- na.omit(roi7)

#tidy roi7
roi7 <- pivot_longer(roi7, cols=2, names_to= "sample", values_to = "ratio")

#create new time variable in minutes
roi7 <- roi7 %>%
  mutate(time_min = timepoint / 60)

roi7$time_min <- as.factor(roi7$time_min)


#graph root2 roi7 10-9M CO4 - panel A
ggplot (roi7, aes(time_min, ratio,   group=1)) +
  geom_line(lwd=1, color = "black") +
  scale_y_continuous(breaks=seq(0.7, 1.2, 0.2), expand = c(0,0), limits= c(0.7,1.2))+
  geom_vline(xintercept="8", linetype = "dashed" )+
  scale_x_discrete(roi7, breaks=c("10", "20", "30"))+
  theme_classic() +
  theme(axis.title.x=element_blank(),
        axis.text = element_text(size = 12),
       axis.title.y = element_blank()
        ) -> plot_roi7_root2

```

###Supplementary Fig S4 - Panel B
```{r}
#select only roi 8 root 3 from the dataset
roi8 <- data[,-(15:20)]
roi8 <- roi8[,-(2:13)]

# rmv NA prior the start of the recording
roi8 <- roi8[-(1:17),] 

#remove NA at the end of the trace
roi8$timepoint <- as.numeric(roi8$timepoint)
roi8 <- roi8%>%
  filter(roi8$timepoint<1835)

roi8 <- na.omit(roi8)

#tidy roi8
roi8 <- pivot_longer(roi8, cols=2, names_to= "sample", values_to = "ratio")

#create new time variable in minutes
roi8 <- roi8 %>%
  mutate(time_min = timepoint / 60)

roi8$time_min <- as.factor(roi8$time_min)


#graph root3 roi8 10-9M CO4 - panel B
ggplot (roi8, aes(time_min, ratio,   group=1)) +
  geom_line(lwd=1, color = "black") +
  scale_y_continuous(breaks=seq(0.7, 1.2, 0.2), expand = c(0,0), limits= c(0.7,1.2))+
  geom_vline(xintercept="8", linetype = "dashed" )+
  scale_x_discrete(roi8, breaks=c("10", "20", "30"))+
  theme_classic() +
  theme(axis.title.x=element_blank(),
        axis.text = element_text(size = 12),
        axis.title.y = element_blank()
        ) -> plot_roi8_root3

```

###Supplementary Fig S4 - Panel C
```{r}
#select root 2 from the dataset
root2 <- data[,-(12:20)]

# rmv NA prior the start of the recording
root2 <- root2[-(1:17),] 

#remove NA at the end of the trace
root2$timepoint <- as.numeric(root2$timepoint)
root2 <- root2%>%
  filter(root2$time<1835)

root2 <- na.omit(root2)

#tidy root2
root2_tidy <- pivot_longer(root2, cols=2:11, names_to= "sample", values_to = "ratio")
root2_tidy <- root2_tidy %>% separate(sample, c('root', 'roi'))

#create new time variable in minutes
root2_tidy <- root2_tidy %>%
  mutate(time_min = timepoint / 60)

root2_tidy$time_min <- as.factor(root2_tidy$time_min)
root2_tidy$roi <- as.factor(root2_tidy$roi)
root2_tidy$root <- as.factor(root2_tidy$root)
root2_tidy$timepoint <- as.factor(root2_tidy$timepoint)


###calculate mean and errors of root by time
root2_stats  <- root2_tidy %>% 
  group_by(time_min) %>% 
  mutate(mean = mean(ratio), 
         stdev = sd(ratio),
         sterr = sd(ratio)/sqrt(length(ratio)),
         UI = paste(time_min, roi, sep = '_')) 

ggplot (root2_stats, aes(time_min, mean,   group=1)) +
  geom_line(lwd=1, color = "#7D7D7B") +
  geom_ribbon(aes(ymin=mean-sterr, ymax=mean+sterr),alpha=0.2,   colour = NA) +  
  scale_y_continuous(breaks=seq(0.8, 1, 0.2), expand = c(0,0), limits= c(0.8,1))+
  geom_vline(xintercept="8", linetype = "dashed" )+
  scale_x_discrete(root2_stats, breaks=c("10", "20", "30"))+
  theme_classic() +
  stat_smooth(color="#FF6A00", size = 1.5, method = "lm", formula = y ~ poly(x, 15))+
  theme(axis.title.x=element_blank(),
        axis.text = element_text(size = 12),
        axis.title.y = element_blank()
        ) -> plot_mean_root2

```

###Supplementary Fig S4 - Panel D
```{r}
#select root 3 from the dataset
root3 <- data[,-(2:11)]

# rmv NA prior the start of the recording
root3 <- root3[-(1:17),] 

#remove NA at the end of the trace
root3$timepoint <- as.numeric(root3$timepoint)
root3 <- root3%>%
  filter(root3$time<1835)

root3 <- na.omit(root3)

#tidy root3
root3_tidy <- pivot_longer(root3, cols=2:10, names_to= "sample", values_to = "ratio")
root3_tidy <- root3_tidy %>% separate(sample, c('root', 'roi'))

#create new time variable in minutes
root3_tidy <- root3_tidy %>%
  mutate(time_min = timepoint / 60)

root3_tidy$time_min <- as.factor(root3_tidy$time_min)
root3_tidy$roi <- as.factor(root3_tidy$roi)
root3_tidy$root <- as.factor(root3_tidy$root)
root3_tidy$timepoint <- as.factor(root3_tidy$timepoint)


###calculate mean and errors of root by time
root3_stats  <- root3_tidy %>% 
  group_by(time_min) %>% 
  mutate(mean = mean(ratio), 
         stdev = sd(ratio),
         sterr = sd(ratio)/sqrt(length(ratio)),
         UI = paste(time_min, roi, sep = '_')) 

ggplot (root3_stats, aes(time_min, mean,   group=1)) +
  geom_line(lwd=1, color = "#7D7D7B") +
  geom_ribbon(aes(ymin=mean-sterr, ymax=mean+sterr),alpha=0.2,   colour = NA) +  
  scale_y_continuous(breaks=seq(0.8, 1, 0.2), expand = c(0,0), limits= c(0.8,1))+
  geom_vline(xintercept="8", linetype = "dashed" )+
  scale_x_discrete(root3_stats, breaks=c("10", "20", "30"))+
  theme_classic() +
  stat_smooth(color="#FF6A00", size = 1.5, method = "lm", formula = y ~ poly(x, 15))+
  theme(axis.title.x=element_blank(),
        axis.text = element_text(size = 12),
        axis.title.y = element_blank()
        ) -> plot_mean_root3

```

###Assemble panels into Supplementary Figure S4
```{r}
plot_roi7_root2+
  plot_roi8_root3+
  plot_mean_root2+
  plot_mean_root3+
  plot_annotation(tag_levels = 'A') +  
  plot_layout(guides = 'collect', heights = c(1, 1, 1.7, 1.7), ncol = 1) ##-> plot_figS4

```
````
