## Supplementary material for "Unlocking the multiphasic nature of intracellular calcium signatures triggered by fungal signals in *Lotus japonicus* roots": Table S1

Table S1. List of primers used in this study

| Name | Sequence 5' --> 3' | Description |
| --- | --- | --- |
| Primers used for cloning | | |
| C58 | AACAGGTCTCAGGCTCCATGGCCAATTGTTGCTCTCACG | aequ_cyt_F_moduleC |
| C59 | AACAGGTCTCACTGAGGGGACAGCTCCACCGTAGAG | aequ_cyt_R_moduleC |
| C96 | AACAGGTCTCAGGCTCCATGTTGCAGCCTAAGAAGAAGAGAAAGGTTGGAG | NLS_YFP_AEQ_F moduleC |
| C34 | AACAGGTCTCAACTACGACGAGTCAGTAATAAACGG | L0.modF_AtUBQ10_promoter_F |
| C35 | AACAGGTCTCAATACAACATACACTTTCACTTGTGATC | L0.modG_AtUBQ10_terminator_R |
| C75 | AACAGGTCTCAACCTGGAGGGAGAGAGGATTTTGAG | pLjUBQ10_modA_F |
| C76 | AACAGGTCTCATGTTCTGTAATCACATCAACAACAGATA | pLjUBQ10_modB_R |
| Primers used for qRT-PCR | | |
| UBI_F | TTCACCTTGTGCTCCGTCTTC | *LjUbiquitin* For (Giovannetti et al., 2015) |
| UBI_R | AACAACAGAACACACAGACAATCC | *LjUbiquitin* Rev (Giovannetti et al., 2015) |
| Q47 | GAAATGGCGTACTGTCTTTAAACCA | *LjRbohB-like* For (Bozsoki et al., 2017) |
| Q48 | GGTGGTTTTCCTGGAAAAGTCAAGA | *LjRbohB-like* Rev (Bozsoki et al., 2017) |
| Q49 | GCAGTGTCTGATTAGTTAGTCTCA | *Ljchitinase* For (Bozsoki et al., 2017) |
| Q50 | GAAGTGATCACATACAGTACTCAAC | *Ljchitinase* Rev (Bozsoki et al., 2017) |
| Q51 | TCTCAATTCCCCAGGAGTTACTTC | *LjWRKY70-like* For (Bozsoki et al., 2017) |
| Q52 | TTGATCGGAGTGTCTTTGCATG | *LjWRKY70-like* Rev (Bozsoki et al., 2017) |
| Q53 | CATGGTTATGATGTTACTGCTCGT | *LjPRp27-like* For (Bozsoki et al., 2017) |
| Q54 | GATCACTATAACCAGTCCTCATCT | *LjPRp27-like* Rev (Bozsoki et al., 2017) |
