## Supplementary material for "Unlocking the multiphasic nature of intracellular calcium signatures triggered by fungal signals in *Lotus japonicus* roots": Table S2

Table S2. List of plasmids used in this study

| Name | Type | Resistance | SOURCE | Description |
|---|---|---|---|---|
| pGGB003 | entry | ampicillin | Addgene - GreenGate cloning system | dummy - module B (N-tag) |
| pGGA006 | entry | ampicillin | Addgene - GreenGate cloning system | Arabidopsis UBQ10 promoter - Module A |
| pGGC015 | entry | ampicillin | Addgene - GreenGate cloning system | mCherry - Module C |
| pGGD002 | entry | ampicillin | Addgene - GreenGate cloning system | dummy - Module B |
| pGGE009 | entry | ampicillin | Addgene - GreenGate cloning system | Arabidopsis UBQ10 terminator - Module E |
| pGGA000 | entry | ampicillin and chloramphenicol | Addgene - GreenGate cloning system | Empty entry vectors (ccdB+ ) - Module A |
| pGGC000 | entry | ampicillin and chloramphenicol | Addgene - GreenGate cloning system | Empty entry vectors (ccdB+ ) - Module C |
| GG59 | entry | ampicillin | this paper | Lotus 2200bp UBQ10 promoter - Module A |
| GG70 | entry | ampicillin | this paper | Selection cassette (pAtUBI_mCherry_tUBI) - Module F |
| GG109 | entry | ampicillin | this paper | CPK17G2A-NES-YFP-linker-aequorin - Module C |
| GG110 | entry | ampicillin | this paper | NLS-YFP-linker-aequorin - Module C |
| | Destination | kanamycin and chloramphenicol | kind gift of Marco Incarbone - GMI, Vienna, Austria | Empty Destination vector - pGGsun_GA |
| GG60 | expression | kanamycin | this paper | CPK17G2A-NES-YFP-linker-aequorin under the control of the LjUBQ10 promoter together with the mCherry transformation marker in pGGsun-GA |
| GG62 | expression | kanamycin | this paper | NLS-linker-aequorin under the control of the LjUBQ10 promoter together with the mCherry transformation marker in pGGsun-GA |
