## Supplementary figures and images for "Unlocking the multiphasic nature of intracellular calcium signatures triggered by fungal signals in *Lotus japonicus* roots"

### Fig_S1_Binci_et_al.jpg

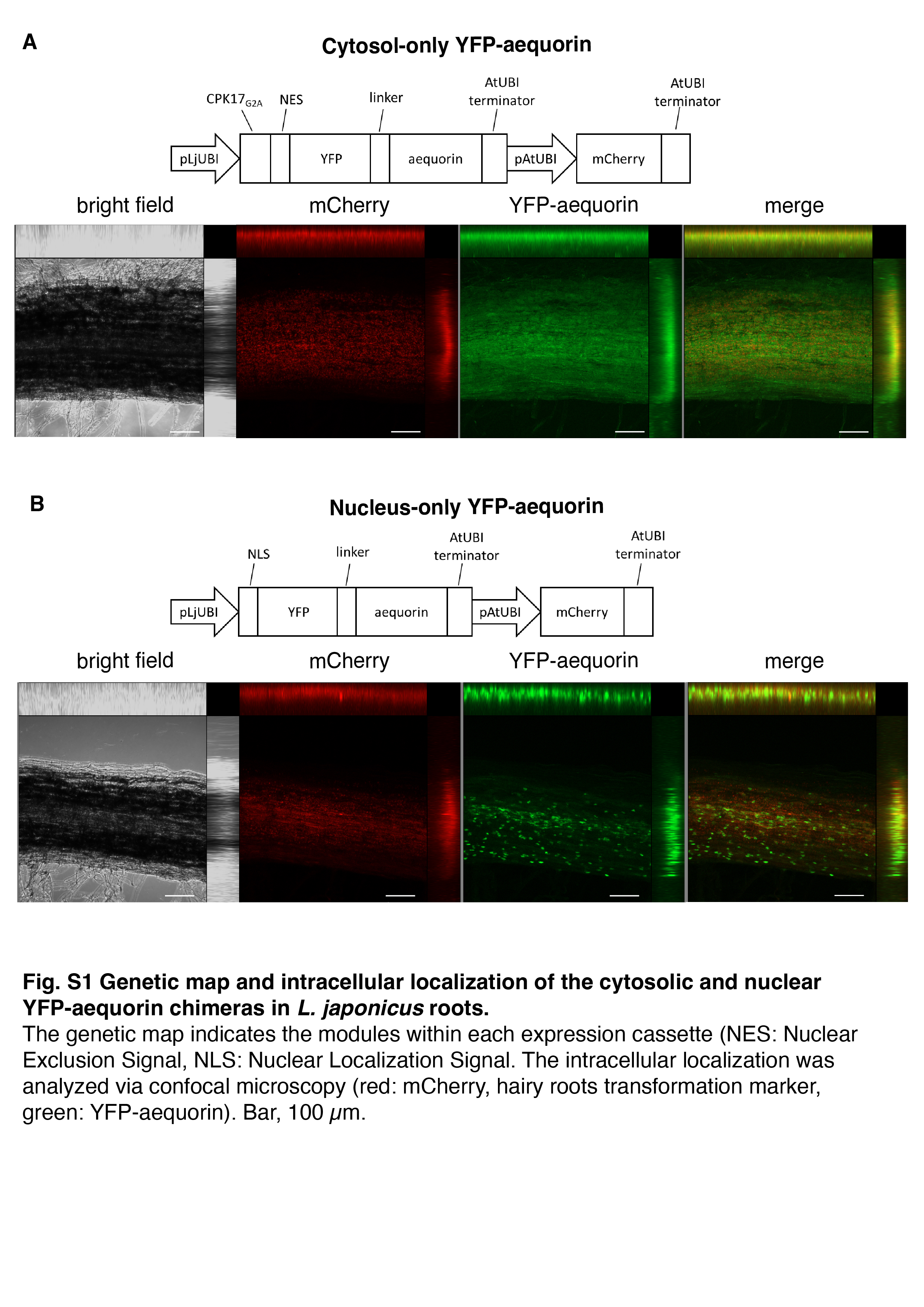
